# KlebPhaCol: A community-driven resource for Klebsiella research identified a novel gut phage order associated with the human gut

**DOI:** 10.1101/2024.12.02.626457

**Authors:** Daniela Rothschild-Rodriguez, Kai S. Lambon, Agnieszka Latka, Ana Rita Costa, Anna Mantzouratou, Claire King, Dimitri Boeckaerts, Elizabeth Sheridan, Francesca Merrick, Francis Drobniewski, Ilaria De Angelis, Kordo Saeed, J. Mark Sutton, Matthew E. Wand, Michael Andrew, Morgen Hedges, Stan J. J. Brouns, Pieter-Jan Haas, Simran Krishnakant Kushwaha, Sophie T. Lawson, Stephen M.E. Fordham, Yan-Jiun Lee, Yi Wu, Yves Briers, Peter R. Weigele, Franklin L. Nobrega

## Abstract

The growing threat of multidrug-resistant *Klebsiella pneumoniae*, coupled with its role in gut colonisation, has intensified the search for new treatments, including bacteriophage therapy. Despite increasing documentation of *Klebsiella-*targeting phages, clinical applications remain limited, with key phage-bacteria interactions still poorly understood. A major obstacle is fragmented access to well-characterised phage-bacteria pairings, restricting the collective advancement of therapeutic and mechanistic insights. To address this gap, we created the Klebsiella Phage Collection (KlebPhaCol), an open-source resource comprising 53 phages and 74 *Klebsiella* isolates, all fully characterised. These phages span five families – including a novel order, *Felixvirales*, associated with the human gut – and target 27 sequence types (including ST258, ST11, ST14) and 28 capsular-locus types (including KL1 and KL2), across six *Klebsiella* species. Freely accessible at www.klebphacol.org, KlebPhaCol invites the scientific community to both use and contribute to this resource, fostering collaborative research and a deeper understanding of *Klebsiella-*phage interactions beyond therapeutic use.

## Introduction

The global rise of antimicrobial resistance (AMR) has prompted urgent action to develop new, effective therapies (1–6), with bacteriophage (phage) therapy emerging as a promising option (7, 8). Phages, as natural predators of bacteria, can precisely target bacterial pathogens, but a reliable pipeline from phage isolation to clinical application remains elusive

(9). Key challenges include limited regulatory frameworks and gaps in understanding phage-bacteria and phage-host interactions, which are essential for developing safe and reliable therapies (10).

*Klebsiella pneumoniae,* a multidrug-resistant pathogen and one of the six “ESKAPE” organisms, exemplifies these challenges. Known for causing severe infections, *K. pneumoniae* has developed resistance to last-resort antimicrobials (11), including carbapenems (12), making it a high priority target for new antimicrobials (13). *K. pneumoniae* infections, including pneumonia, sepsis, and liver abscess, are often acquired in hospital settings but are also found in community-acquired cases, especially involving hypervirulent strains (14–20). The role of this pathogen in chronic gut colonization has further implicated it in gut conditions like inflammatory bowel disease (IBD) (21, 22) and primary sclerosing cholangitis (23), establishing *K. pneumoniae* as a significant gut-associated pathobiont.

A major challenge in phage therapy against *K. pneumoniae* is its highly variable capsule antigen (K-types), with over 180 distinct types now genomically identified (24–27) and associated with different species (28) and virulence traits (29). The diversity complicates treatment because capsule-specific phages, which depend on capsule polysaccharides to bind and infect cells, often have limited host ranges (30–32). While some phages can bind alternative receptors like the O-antigen (33), capsule diversity remains a critical barrier. Beyond receptor diversity, bacterial defence systems and mobile genetic elements can further restrict phage efficacy (34–37). These multifaceted interactions highlight the need for well-characterised phage collections, which can enable researchers to systematically study and address obstacles to successful therapy. Several collections of *Klebsiella* phages have been reported in the literature (33, 34, 38–42). While these mark milestones in the field, there remains a fundamental need for centralising and standardising resources to make them easily accessible for the academic and clinical communities. Standardised, referenced collections, such as the BASEL collection for *Escherichia coli* phages, demonstrate how accessible resources can foster shared advancements (43, 44). Addressing this need, we present the <u>Kleb</u>siella <u>Pha</u>ge <u>Col</u>lection (KlebPhaCol), an open-source collection that contains 53 diverse phages and 74 *Klebsiella* strains, each comprehensively characterised. The open-source nature of KlebPhaCol (available at klebphacol.org) invites the scientific and medical community to contribute additional isolates and data, fostering and evolving this community-driven platform. In addition to informing phage therapy, this collection can be utilised to study fundamental aspects of phage-bacteria interactions. By centralizing and sharing these resources, KlebPhaCol aims to bridge current gaps, empowering the scientific community to collectively advance research on *Klebsiella* and its phages for both therapeutic and broader biological insights.

## Material and methods

### Phage isolation and purification

Numbered phages (e.g. Roth01) were sourced from hospital wastewater effluent collected at the University Medical Centre Utrecht in the Netherlands in 2020 as previously described (45), while lettered phages (e.g. RothC) were sourced from effluent collected at Portswood in Southampton, United Kingdom in 2021. Thirty-two isolates with clinically relevant ST-types were used as isolation hosts (Supplementary Table S2). Based on ST-type grouping, 7 enrichment cultures were produced: 1) ST11 (n=5), 2) ST101 (n=5), 3) ST15 (n=4), 4) ST258 (n=5), 5) ST14 (n=6), 6) ST323 (n=2), and 7) the remaining ST-types (ST489 (n=1), ST86 (n=1), ST38 (n=1), ST23 (n=2)). 50 µL of each overnight culture grown in Lysogeny Broth (LB; Formedium LB-Broth Lennox) were added to each respective enrichment containing 50 mL of LB and 50 µL of the phage source filtrate. Enrichments were incubated overnight at 37°C and shaking at 180 rpm. These were then centrifuged (8, 000 × *g*, 20 min, 4 °C), filter-sterilized (0.45/0.22 µm PES) and kept at 4 °C for subsequent assays. 5 µL of the resulting supernatants were spot-tested for the detection of phage against all 32 isolates using a double-layer agar technique (top agar 0.6%) (46). Susceptible isolates were subsequently plated with serially diluted phages to identify distinct plaque morphologies, which were then single picked with sterile toothpicks, dotted, and spread with sterile paper onto fresh bacterial lawns to purify the phages. This latter step was repeated twice to obtain a consistent plaque morphology. Individually purified phages were then propagated in LB media with their respective host, centrifuged, and filter-sterilized (0.45 µm PES, 0.22 µm PES for final batch productions). The resulting 53 phage stocks were stored at 4 °C and used for further characterisation.

### Transmission electron microscopy

One mL of each phage lysate was sedimented at 21, 0001× *g* for 11h, the top 900 µL were discarded and the remaining volume was resuspended in 900 µL of sterile MilliQ water. The centrifugation and washing steps were repeated once more. Concentrated phages (5 µL) were deposited and incubated for 30 sec on glow-discharged carbon-coated type-B 400 mesh grids (Ted Pella). Excess phage was dried with filter paper before staining with 5% Ammonium Molybdate (w/v) and 0.1-1% Trehalose for 10 sec. Samples were examined in a transmission electron microscope (TEM FEI Tecnai T12) at an acceleration voltage of 80 or 120 kV, and phage particles were examined at 16, 500–105, 000 x magnification. Fiji software (47) v2.9.0/2.14.0 was used to measure the phage lengths and to crop the images to scale.

### Phage host-range

Ten-fold serial dilutions of the phages were spotted onto double-layer agar plates of a total of 74 *Klebsiella spp.* strains. The plates were incubated overnight at 37 °C and phage plaques were observed to distinguish between productive infection (lysis with individual plaques), no infection (lack of plaques) and undetermined lysis (opaque lysis without individual plaques). Assays were conducted in both LB and Tryptic Soy Broth (TSB; Hach Bacto^TM^ Tryptic Soy Broth) media. Unless otherwise stated, all other phage assays were done in LB.

### One-step growth curves

One respective phage per phylogenetic clade was chosen at random to cover the range of phage diversity in the collection. Overnight cultures of the isolation strains were diluted 1:100 in LB and incubated at 37 °C at 180 rpm up to an optical density (OD_600_) of 0.4-0.5. 10 mL of culture were then centrifuged at 4-7, 000 x g for 5 min and the cell pellet resuspended in 5 mL of LB. Five mL of phage [10^5^ PFU/mL] was then added and let adsorb for 5 min at 37 °C at 180 rpm. The centrifugation step was repeated, and the cell pellet resuspended in 10 mL of LB, transferred into a conical flask, and incubated at 37 °C at 180 rpm for 40-120 min. Samples were taken at time 0 and at 5-min intervals during the first 50 min and 10-min intervals thereafter. Samples were immediately 10-fold serially diluted and plated with the isolation strain for plaque quantification in triplicate.

### Plate reader liquid assays

Overnight bacterial cultures of strains susceptible to ST323-phages on solid agar were diluted 1:100 in LB and incubated at 37 °C at 180 rpm up to an OD_600_ of ∼0.3. Cultures were normalized to an OD_600_ of 0.1 and dispensed into a 96-well plate. Experimental wells had phage added at the desired high (≥1) or low (≤1) MOI. Growth was monitored every 10 min for 900 min in a Spectrostar Nano (BMG Labtech) plate reader at 37 °C, non-shaking, in either aerobic or anaerobic conditions. To ensure the latter, all holes in the plate reader were plugged as specified by the manufacturer, and N_2_ gas was consistently pumped at a low rate to eliminate any oxygen for the entirety of the experiment. Growth curves were converted to area under the curve using GraphPad prism and centroid index was calculated by inputting the average OD_600_ values of the three repeats into the Centroid Index Calculator software as per Hosseini *et al* (48).

### Bacteriophage insensitive mutants

BIMs of capsule-deficient strain 51851 were obtained after spot tests with different phages. A random selection of 21 BIMs from 51851 strain were cultured for sequencing and for phage re-testing. Strain DNA was extracted and sequenced as described below to confirm mutations. Phage susceptibility re-testing was performed in a 96 well plate by growth in LB broth at 37°C of the BIMs with the addition of phage. Growth of the bacterial isolate in the absence of phage was used as a positive control. OD_600_ readings were taken every hour up to 20 hrs using a CLARIOstar Plus plate reader (BMG Labtech). Growth curves were analysed and, where there was no observable difference in the presence and absence of phage, this was classified as resistant (i.e. no lysis).

### Phage sequencing, assembly, and annotation

Phage DNA was extracted using phenol-chloroform as previously described (49). DNA from 32 phages were sequenced by BMKGene. For this, sequencing libraries were prepared by BMKGene (Germany) using the Reseq-M DNA kit and paired-end reads (2×150 bp) were generated in the Illumina Novaseq 6000 platform. Approximately 3-4 Gb of clean sequencing data were produced and delivered for each sample, with sequencing depth >5, 000 x. The remaining DNA was sequenced by the UKHSA-GSDU (UK health security agency Genomic Services and Development Unit) (see Supplementary Table S1). Libraries were prepared using the nextera DNA flex library prep kit according to manufactures instructions and reads (2×150 bp) were generated in the Illumina HiSeq 2500 platform. A minimum of 150 Mb of Q30 quality data was obtained for each sample.

Unless otherwise stated, CLC Genomics Workbench v23.0.1 (Qiagen) was used for quality checks, sequence trimming (quality limit = 0.05) and genome assembly. Reads were subsampled then assembled with the *de novo* assembler tool (default parameters) on CLC. Sequencing reads for 14 phages (see Supplementary Table S1) were checked for quality using FastP (50) v.0.12.4 and Soapnuke (51) v2.1.7 with default parameters. For these specified phages, reads were sampled and trimmed using Seqtk v1.3.0 and then assembled using SPAdes (52) v3.13.0. All produced assemblies were manually inspected on Bandage v0.8.1 and Geneious Prime v11.0.18+10.

The phages’ closest relative was determined as the top hit according to the maximum score provided by BLASTn (March-June 2023 and February 2024). Assemblies were mapped to fastq reads to check for irregularities using Qualimap2 v2.3 (53).

Phages were zeroed to allow phage comparisons with canonical phages whereby a conserved feature was chosen to serve as gp1 or “start-site” for each of the families represented in the collection. These genome start-sites were chosen based on historical precedent and/or biology of infection and/or DNA packaging. For the *Straboviridae*, which includes the well-known *Escherichia* phage T4 (genus *Tequatrovirus*), and the genera *Jiadodavirus* and *Slopekvirus*, the *rIIA* gene was chosen, in accordance with NCBI record NC_000866.4 (54). In cases where a landmark feature overlapped another gene, the nearest non-CDS region 5’ or 3’ to rIIA was chosen to avoid software artefacts. The *Demercviridae* contains a landmark member, *Escherichia* phage T5 (genus *Tequintavirus*, NC_005859.1) where the first-step-transfer region encoding dmp, a 5’-deoxyribonucleotidase, is first to enter the cell upon infection (55, 56). The *Drexlerviridae* includes phage T1 (genus *Tunavirus*, NC_005833.1), which is known to have terminal repeats at the genome ends (57). For the *Autographiviridae* Roth32, infection by coliphage T7 (NC_001604.1), a member of the genus *Teseptimavirus* of this family, an ∼850 bp segment of the virion DNA enters the cell first (58). This region is rich in host RNA polymerase promoters and consequent gene transcription drives the entry of the remaining genome into the cell (58). Phages from the *Drulisvirus* genus were zeroed to the terminase subunit, a convention built into some automated annotation pipelines (59). The novel *Felixviridae* family was zeroed for the core region to come first. Manual assignment of nucleotide start-site was accomplished using Geneious Prime v11.0.18+10.

Final phage genome length and GC content were determined by EMBOSS v6.6.0.0. Phage sequences were then inputted to PhageTerm (60) via the Center for Phage Technology galaxy portal (https://cpt.tamu.edu/galaxy-pub), to identify phage termini and packaging.

Phage coding sequences (CDS) were predicted with PHANOTATE (61) v2019.08.09 using translation table 11, then annotated using multiPHATE (62) v2.0.2 against the NCBI database selecting annotations with an Evalue threshold of 0.001. tRNA genes were identified using tRNAscan-SE (63) v2.0.12 via multiPHATE and confirmed using ARAGORN (64) v1.2.41, although tRNAscan-SE findings were kept. *Sugarlandvirus* had two predicted tRNAs that overlapped with hypothetical protein CDSs, and were therefore removed for the GenBank submission. Phages were also annotated with the Pharokka (59), Phold (65), and Domainator (66) v0.7 to highlight additional domain and gene functions (Domainator annotations are available on the Figshare version of the phages). Annotations were manually curated using Geneious Prime v11.0.18+10. Anti-defence proteins were predicted using AntiDefenseFinder (67). Potential antimicrobial resistance and virulence genes in the phages were predicted using the CARD database and VFDB, respectively.

The lifestyle of the phages was predicted using Bacphlip (68). Phage receptor-binding proteins (RBPs) and depolymerases were identified using RBPdetect (69) v3.0.0. and DepoScope (70) v1.0.0, respectively. Typical tailspike folds were verified with AlphaFold v2.3.1 (71) modelling. The selection was manually curated for depolymerases based on conserved structural elements according to the method described previously (72). When a RBP is not identified as a depolymerase, the RBP is presumed to be a tail fibre that binds to the bacterial receptor.

### Phage DNA-modifying enzymes

Phages encoding DNA-modifying enzymes were manually curated and annotated. Their nucleoside composition was analyzed using high-performance liquid chromatography coupled mass spectrometry (HPLC-MS) on1enzymatically hydrolyzed DNA. Between 1-2 µg of extracted phage DNA (by phenol-chloroform as described above) was treated with Nucleoside Digestion Mix (New England Biolabs, #M0649) following the manufacturer’s protocol at 37°C overnight. The nucleoside mixture1was filtered1through a PTFE 0.2 µm centrifugal filter, and the filtrate1was subjected1to the reverse phase HPLC-MS for nucleoside separation and detection. LC-MS instrumentation was performed on an Agilent 1290 Infinity II UHPLC-MS system equipped with a G7117 Diode Array Detector and an LC/MSD XT G6135 Single Quadrupole Mass Detector. A Waters XSelect HSS T3 C18 column (2.5 µm, 2.1 mm × 100 mm) was used for the chromatography and operated1at a flow rate of 0.6 mL/min with a binary gradient mobile phase consisting of 10 mM ammonium acetate (pH 4.5) and methanol.1The course of chromatography was monitored by UV absorbance of the effluent1at 260 nm. Mass spectrometry1was operated1in both1positive (+ESI) and negative (-ESI) electrospray ionization modes. MS1was performed1with a capillary voltage of 2500 V at both ESI modes, a1fragmentor1voltage of 70 V, and a mass range of m/z from 100 to 1000.1Agilent ChemStation software was used to process primary LC-MS data. Adobe Illustrator was used to compile, render, and annotate the chromatograms exported from ChemStation software.

### Phage comparative genomics

To assign phage taxonomy, genomes were run on PhageGCN (73) web server and confirmed by clustering on vContact2 (74) v0.11.3, using the default database and visualised using Cytoscape v3.10.2. Intergenomic similarity was calculated using VIRIDIC (75) on the web server and similarity matrices were re-plotted using Pheatmap (76) v1.0.12. Phylogenetic analyses were produced by the VICTOR web server with default settings, which employs the Genome-BLAST Distance Phylogeny method adapted to bacteriophages (77). Tree images were rendered and rooted at the midpoint using iTOL (https://itol.embl.de/) (78) v6.1.1. Synteny plots were produced by Clinker (79) on their web server.

### Taxonomic classification of RothC and RothD

To identify the closest relatives to RothC and RothD, a BLASTn search to the NCBI database filtered for the viral taxa ID (10239) was conducted. The top hit with a complete genome (vB_Kpn_Chronis, accession MN013086.1) (80) was compared using Clinker and VIRIDC. To confirm the gut-relevance of phages RothC and RothD, the publicly available Gut Phage Database (GPD) (81) was downloaded and searched via command-line BLASTn v2.15.0 with default settings. Hits were analysed by retrieving their metadata from the GPD, and only high-quality genomes with a mean ± SD genome completeness of 98.3% ± 2.6 were retained for taxonomic enquiry (n=132). To infer order and family rank we employed VipTree (82) via the webserver selecting for all other dsDNA prokaryotic viruses within their database. The sister clades surrounding RothC and RothD were extracted for visualisation but only the family containing RothC and RothD was selected for taxonomic classification below order level. To infer ranks below family-level a demarcation criterion according to ICTV guidelines was applied as follows. Based on complete genome nucleotide comparisons using VIRIDIC and VICTOR tools, ≥45% nucleotide identity to RothC and RothD was used for assigning at the subfamily level; ≥70% nucleotide identity was used for assigning at the genus level; and ≥95% was set for same species classification. The taxonomic proposal was submitted to the ICTV on the 27^th^ of June 2024.

### Bacterial DNA extraction and genome assembly

Seventy-four clinical isolates of *Klebsiella* spp. were used in this study. Sixty-seven are *K. pneumoniae*, two *K. oxytoca,* two *K. variicola,* one *K. aerogenes*, one *K. pneumoniae subsp. ozaenae*, and one *K. similipneumoniae* ; see Supplementary Table S2 for isolate characteristics. Thirty-two strains were used for phage isolation enrichment cultures, but only 18 continued as isolation hosts (Supplementary Table S2).

Genomic DNA for the *Klebsiella* strains were extracted using the GeneJet Genomic DNA kit (ThermoScientific) or the Promega Wizard DNA extraction kit according to the manufacturer’s instructions. DNA was quantified by a Qubit fluorometer using the high sensitivity dsDNA kit (Invitrogen) and Nanodrop. DNA was prepped and sequenced by UKHSA-GSDU as described above. Fastq reads were quality trimmed using Trimmomatic (83) v0.39 and draft chromosome contigs were assembled using SPAdes v3.15.3 filtering out contigs <1kp.

### Bacterial genome analyses

Genomes were annotated using Prokka v1.14.6 (84). Strains were classified by their sequence (ST) and capsular locus (K) types using the multilocus sequence typing (MLST) database (Center for Genomic Epidemiology (https://cge.cbs.dtu.dk/services/MLST/), and Kaptive (85) v2.0.7 using the K locus primary reference database, respectively. Strains from the *K. pneumoniae* species complex (KpSC) (19) were also classified by the cgMLST-based Life Identification Numbers (cgLIN codes) available via Pathogenwatch (https://pathogen.watch/) (86) to provide a better phylogenetic resolution and precision at a nomenclature-based level (87) (Supplementary Table S2). Strains were run through the Kleborate (25) pipeline to obtain virulence and resistance scores, and outputs were visualised using the Kleborate-Viz platform online (https://kleborate.erc.monash.edu/) (no markers were found for strain 163575R). The phylogeny of the strains was calculated via PopPUNK (88) v2.5.0 using the default fitted model for *Klebsiella pneumoniae*. The tree was rendered in iTOL v6.1.1.

The bacterial virulence factors, antibiotic resistance, and stress resistance genes were identified using Abricate (89) v1.0.1 against the Comprehensive Antibiotic Resistance Database (CARD) (90), NCBI AMRFinderPlus (91) and Virulence Factor Database (VFDB) (92). Prophage regions were identified using Phigaro (93) v2.2.6 on default mode. The defence systems in the genomes were identified using PADLOC (94) v1.1.0 and DefenseFinder (95) v1.0.9. Incomplete defence systems, systems VSPR and PDC were removed from quantification analyses but are all included in Supplementary Table S2. Correlation analyses between encoding defence systems and host range outcomes were conducted with Spearman’s correlation and plotted in RStudio v2024.04.2 using the ggplot2 (96) package.

Bacterial capsule locus (defined as the genetic region from *galF* to *uge*) (97) were manually assembled for isolation and production hosts as well as for those highly susceptible strains (total of 22 strains). Assembly was conducted by first looking for the more conserved regions of *galF* and *uge* genes and then individually checking and annotating other genes in Seq Builder v14.0.0 (DNAstar Lasergene). In some cases, due to transposon insertions within the CPS locus, it was not possible to generate one contig containing the complete locus; for such cases a string of n’s was artificially added to represent a break in the contigs.

### Antibiotic susceptibility

For clinical isolates obtained at the University Medical Centre Utrecht (see Supplementary Table S2), antibiotic susceptibility was determined as previously described (45). For the remaining clinical isolates, testing was done by UKHSA as follows. The minimal inhibitory concentrations for antibiotics and biocides were determined by a standard broth microdilution method at a starting inoculum of 5×10^5^ CFU/mL, using Phoenix M50 system (BD Biosciences) and EUCAST breakpoints, with the exception that 96-well polypropylene plates (Griener Bio-One, Ltd.) were used instead of polystyrene plates to test colistin. Plates were scored by eye, looking for no visual growth and confirmed by OD_600_ measurement after 16–20 h with a 0.1 OD_600_ threshold using a CLARIOstar Plus plate reader (BMG Labtech).

### Genomic comparisons of Felixvirales phages

RothC and RothD were taken as the representative phages of the *Felixviridae* family. To determine the prevalence of *Felixviridae* phages within bacterial genomes given their temperate lifestyle, the BV-BRC (https://www.bv-brc.org/) bacterial strain database (n=64, 364; 25, 384 complete high-quality *Klebsiella* spp. and 38, 980 complete high-quality non-*Klebsiella* bacterial genomes), and associated metadata was retrieved (accessed July 2024). The conserved region and the full genomes of RothC and RothD were independently searched against the downloaded bacterial genomes using command line BLASTn v2.15.0 with an Evalue threshold of 0.005 with a -max_target_seqs parameter of 100, 000. All hits were extracted and searched for prophage regions using Phigaro (93) v2.4.0 with default settings. The ten upstream and ten downstream genes from the hit region were extracted for analysis. Significant hits were defined by a hit length ≥ 1000 bp and with an Evalue <10^−8^.

To investigate the predominance of *Felixvirales* in the average human gut, we first looked at the GPD hits used in the taxonomic characterisation of RothC and RothD (see above) and matched them with the GPD’s available metadata. The produced dataset was then analysed. Taxonomic characterisation of RothC and RothD revealed several relatives assembled from a singular study by Tisza *et al* . (98). Thus, we gathered all returned hits from the online BLASTn server and matched accession queries to those coming from the mentioned study. This resulted in a total of 229/406 total hits matching their chronic disease dataset, of which 205/229 (90%) were high-confidence hits with an Evalue ≤10^−8^. We then matched these to the study’s metadata and analysed the resulting dataset.

The relative abundance of *Felixvirales* phages was calculated as follows. The quality filtered reads from a subset of 117 ‘healthy’ human stool metagenomes from the Human Microbiome Project (99, 100) were retrieved and aligned to a set of 133 *Felixvirales* phage genomes using the end-to-end alignment mode of Bowtie2 v2.5.4 (101). Bacterial reads were identified using Kraken2 v2.1.3 (102). A count table of reads aligned to contigs and total number of reads per metagenome was generated with Samtools v1.20 (103) and imported into Rstudio v2024.04.2+764 for analysis. Packages ggplot2 (96) and ggbreak (104) were used for plots.

### PCR detection

Primers were designed to target gp7, a hypothetical protein (or putative virion structural protein by Phold) that maintains a high conservation across the *Felixviridae*: Forward 5’-ATGTTCCGTCAGGGGAAGTTC-3’, Reverse 5’-AAGCCTGGTTGTTAAAACTGG-3’. Primers were synthesized by IDT. Reactions were done with OneTaq quick load (NEB M0486) according to the manufacturer’s instructions on a Bio-Rad T100 Thermal Cycler and visualised on a 0.7% agarose gel. Positive controls were RothC and RothD. Negative control was prepared using DNase-free water instead of template DNA. Specificity to *Felixviridae* phages was confirmed by also testing phages T4 (as a non-*Klebsiella* phage control) and Roth32 (as a *Klebsiella* phage control). The presence of *Felixviridae* phages in the environment was also tested by using filtered raw effluent from sewage plants in Southampton and Petersfield as well as ocean water from the Isle of Wight, UK (collected in the summer of 2024), filtered through a Vivaflow® 200 cassette recirculation system (Sartorius) and then through a 0.45 µm PES membrane. All controls (except for the negative control) were first heated at 95 °C for 5 min to break virion capsules before adding as template DNA to the reactions. PCR products were cleaned and concentrated with the ThermoFisher GeneJET PCR purification kit and sent for sequencing at Plasmidsaurus UK. Reads were trimmed and quality filtered using FastP v0.12.4 (50) on the fastplong parameter, and then mapped to RothD_gp7 (100% identity to RothC_gp7) using minimap2 v2.28-r1209 (105). Coverage depth was obtained with Samtools v1.20 (103) and Bedtools v2.30 (106) and results were imported in table format to RStudio v2024.04.2+764 and plotted using the ggplot2 (96) package.

## Results

### Overview of KlebPhaCol

KlebPhaCol is an open-source *Klebsiella* phage and strain collection for the research community, which includes the biological material (phages and strains), as well as the accompanying data. The goal of this collection is to provide easy, cost-effective access to phages and bacterial strains for collaborative research exploring interactions between phages, bacteria, and humans in the context of *Klebsiella* infections, ultimately contributing to phage therapy developments by centralising knowledge on representative sets of phages and strains. To facilitate accessibility, all information and data about the collection has been collated into a single domain online, www.klebphacol.org. The collection and its associated data can be searched on this website, and users (from the academic or medical community) can also submit their requests for access to the collection via a simple form in the same domain (subject to verification). Given the large repertoire of *Klebsiella* phages reported in the literature, with no easy access to them, we open KlebPhaCol to be expandable, where researchers can deposit their *Klebsiella* phages and strains to be shared openly alongside its current contents. All data produced using the collection will be made available via the website too, allowing users to find the published collaborations in one place. KlebPhaCol went live on August 2023 and the resource has been shared with 21 labs from eight countries since.

KlebPhaCol comprises phages isolated using 32 clinically relevant *Klebsiella spp.* strains provided by the UK Health Security Agency (UKHSA), with sewage filtrates from two countries serving as phage sources (Figure 1A, Supplementary Tables S1 and S2). A total of 53 phages were successfully sequenced and characterised for inclusion in the collection (Figure 1B). These phages represent five families, including a proposed new family, span seven genera, and encompass the three main morphotypes of double stranded DNA (dsDNA) tailed phages (Figure 1C). All but two phages are strictly virulent (Figure 1B). The phage genomes range in size from 43, 182 to 174, 596 base pairs (Figure 1B, Supplementary Table S1).

**Figure 1.**
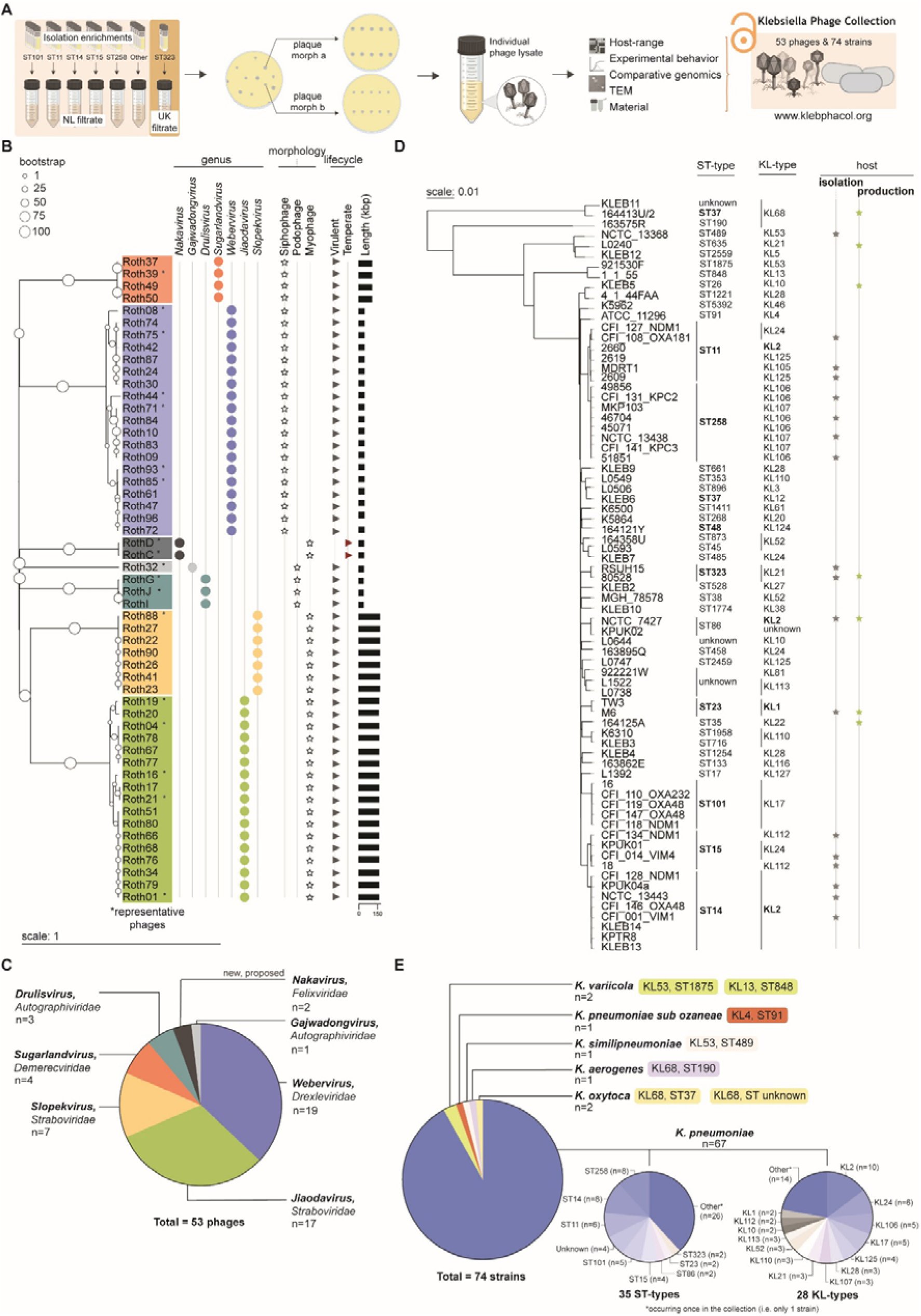
Overview of the Klebsiella Phage Collection (KlebPhaCol). (**A**) Phage isolation enrichment groups and methods that resulted in the phage collection of KlebPhaCol (NL sewage filtrate originated from University Medical Center, Utrecht, Netherlands; UK sewage filtrate originated from Portswood, Southampton, United Kingdom). Figure created with elements from BioRender, edited in Adobe illustrator. (**B**) Phylogeny of the 53 phages of the collection and associated data. Phylogenetic tree was calculated using a Genome BLAST Distance Phylogeny method, and midpoint rooted. (**C**) Quantification of the phage taxa covered by the phages in KlebPhaCol. (**D**) Phylogeny of the 74 strains of the collection and associated data. Phylogenetic tree was produced by PopPUNK and midpoint rooted. (**E**) Quantification of the species of *Klebsiella,* their sequence type (ST) and capsule locus type (KL), included in KlebPhaCol. All trees were rendered in iTOL.

To facilitate reproducibility and shareability, we selected seven strains as production hosts for the entire collection (Figure 1D). Currently, the collection is composed of 74 strains, of which 69 are clinical isolates from different countries (Supplementary Table S2), while the remaining five are ATCC/NCTC-type strains. These 74 strains represent six *Klebsiella* species, 40 sequence types (STs), 32 capsule locus (KL) types, and 10 O-antigen (O) types (Figure 1E, Supplementary Table S2).

### KlebPhaCol phage characterization

The characterisation of KlebPhaCol phages involved genomic, phenotypic, and behavioural analyses, which we organised by genus (Figure 2, Supplementary Figures S1 and S2).

**Figure 2.**
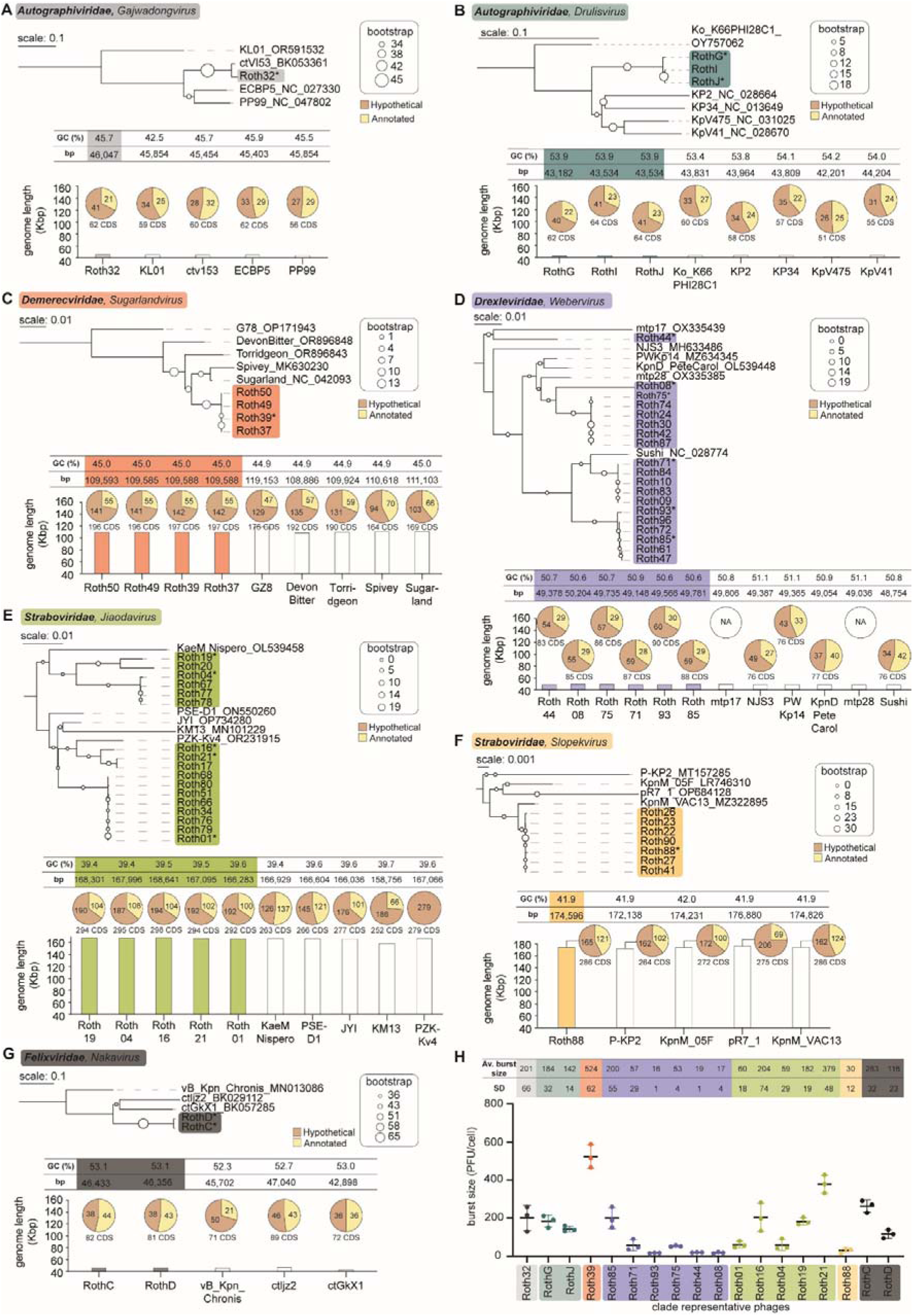
Comparisons between representative phages of KlebPhaCol and their close relatives. (**A-G**) Panels contain a genome-based phylogenetic tree of the representative phages and close relatives for each genus. All trees were calculated by VICTOR and rooted at the midpoint, visualised by iTOL. The tables display the GC content and base pair (bp) length of each genome. The latter is also visualized in the bar graphs relative to the largest genomes of the collection (>160 Kbp). The pie charts illustrate the proportion of annotated coding sequences (CDS) versus those annotated as ‘hypothetical protein’ in this study for the Roth phages, and as deposited in GenBank for the close relatives. ‘NA’ indicates the deposited genome was not annotated. (**H**) Burst sizes of the representative Roth phages calculated by one-step growth curve experiments conducted in triplicate for each phage. The average (av) and standard deviation (SD) burst size of each is also given in table format.

Genomic characterisation focused on comparisons with existing related phages, incorporating phylogenetic, synteny, and gene content analyses. Receptor-binding proteins (RBPs) and depolymerases were predicted for each phage, highlighting key proteins involved in receptor recognition (Supplementary Table S3). DNA-modifying genes were also identified and their corresponding DNA modifications were experimentally verified by HPLC (Supplementary Figures S2 and S3, Supplementary Table S1).

Phenotypic analysis included plaque morphology and TEM imaging (Supplementary Figure S4), while behavioural analyses involved one-step growth curves (Supplementary Figure S5) and host-range determination (Supplementary Figures S6 and S7). The subsequent sections provide detailed descriptions of the characteristics of these phages, organised by genus. Representative phages were selected from groups that displayed high intergenomic similarity (Supplementary Figure S1A).

#### Gajwadongvirus

Roth32 is the only *Gajwadongvirus* of the *Autographiviridae* family in KlebPhaCol. This podophage (Supplementary Figure S4A) has a genome size of 46, 047 bp (Figure 2A) and a burst size of 201±66 particles per lytic cycle (Figure 2H, Supplementary Figure S5A). Its closest relative was found to be a metagenome-assembled phage with a partial genome, ctVI53 (Genbank: BK053361, Figure 2A). Roth32 has 62 coding sequences (CDS), of which only 21 are functionally annotated (Figure 2A). Genome synteny is not well maintained with its relatives, apart from around 29/62 of its CDS, suggesting its uniqueness (Supplementary Figure S2A).

Roth32 encodes one of the only four predicted depolymerase enzymes within phages of the collection, Roth32_gp7. This enzyme is homologous to a lyase (PHYRE2: c4y9vA, confidence 80.2, coverage 58%, aa 128-515) and a hydrolase (PHYRE2: c3eqnB, confidence 93.6, coverage 49%, aa 258-576), which is characteristic for depolymerases. Roth32_gp7 shares 57.36%, 57.67% and 57.21% identity on the amino acid level with the depolymerases of the *Klebsiella* phages NTUH-K2044-K1-1 (Genbank: YP_009098385), KpV71 (Genbank: YP_009302756) and GBH001 (Genbank: GBH001_056), respectively, which were all described to be K1-specific (107–109). In agreement with this, Roth32 infects only strain M6, a KL1 strain (Supplementary Table S2, Supplementary Figure S6). Overall, these data suggest that Roth32_gp7 may be a K1-specific depolymerase.

Roth32 had one additional RBP predicted, Roth32_gp59. PHYRE2 (110) analyses showed that it is homologous to the N-terminus of the bacteriophage T7 tail fibre protein gp17 (PHYRE2: c7bozj, confidence 100%, coverage 46%, aa 4-163), which functions as the anchor of the tail fibre to the T7 tail, suggesting that Roth32_gp59 may directly interact with the Roth32 tail. Based on high probability alignments to the tailspike protein 4 (TSP4) of phage CBA120 (Genbank: NC_016570.1) via HHPred, we also found that Roth32_gp59 is equipped with a T4 gp10-like branching domain which is a putative docking site for other RBPs, where the branching domain sequence is embedded in the 163-306 aa of Roth32_gp59. Therefore, Roth32_gp59 could be an intermediate adaptor protein to which another true RBP can bind (111). These data suggest that the RBP architecture of Roth32 is similar to phages from the previously described KP34 viruses group A (72), where RBP1 (gp59 in Roth32) is truncated and anchors the RBP system as an adaptor protein to the phage tail and provides a branching site for RBP2 (gp7 in Roth32) with depolymerising activity.

#### Drulisvirus

Phages RothG, RothI, and RothJ belong to the *Drulisvirus* genus of the *Autographiviridae* family (Figure 2B). These podophages (Supplementary Figure S4A) have ≈43 kb genomes (Figure 2B) of high similarity (>99%; Supplementary Figure S1A) and infect the same isolation host, RSUH15, an ST323 KL21 *K. pneumoniae* strain. Phage vB_Ko_K66PH128C1 (Genbank: OY757062) is their closest relative at the time of our search (Figure 2b). RothI and RothJ are more closely related to each other than to RothG. As a result, we selected RothG and RothJ as representatives of this genus for further characterisation. RothG and RothJ have an average burst size of 182±32 and 142±14 particles (Figure 2H), encoding 62 and 64 CDS of which only 22 and 23, respectively, are annotated (Figure 2B). Genome synteny is well-kept with their relatives except for a highly variable region with little to no identity between the related phages from gp9-gp15 of RothG, RothJ and RothI (Supplementary Figure S2B).

Besides Roth32, RothG, RothI and RothJ are the only phages of the collection encoding RBPs with depolymerase activity (based on high-confidence predictions and manual curations): RothG_gp8, RothI_gp8, and RothJ_gp8. These depolymerases share a high aa identity of 98.43%. Their gp8 shares similarity with KL21-targeting proteins of other *Klebsiella* phages (YP_010843553 – 36.83% identity, YP_009198669 – 35.28% identity, YP_003347556 – 35.28% identity to RothG_gp8) (32, 112, 113), suggesting that RothG_gp8, RothI_gp8, and RothJ_gp8 may be KL21-specific depolymerases. Indeed, these phages infect KL21 strains of the collection, but they additionally infect KL4 and KL53 strains (Supplementary Figure S6). It is possible that gp8 also facilitates KL4 and KL53 targeting, or that the phages encode additional unknown depolymerases that facilitate this expanded host range (Supplementary Table S3).

Like Roth32, RothG, RothI and RothJ also encode an additional RBP homologous to the N-terminus of the bacteriophage T7 tail fibre protein gp17, RothG_gp62, RothI_gp63, and RothJ_gp64. High probability HHPred analyses show the branching domain sequences are embedded in the 191-259 aa of RothG_gp62, RothI_gp63, and RothJ_gp64. As for Roth32_gp59, this suggests that these RBPs are intermediate adaptor proteins to which another RBP can bind, and that the RBP architecture of phages RothG, RothI and RothJ is the same as Roth32 and therefore similar to phages from the previously described KP34 viruses group A (72). The depolymerising activity of the four podophages of the collection is also confirmed by the haloed plaques produced by these phages (Supplementary Figure S4A).

#### Sugarlandvirus

KlebPhaCol contains four phages of the *Sugarlandvirus* genus (*Demerecviridae* family), Roth37, Roth39, Roth49 and Roth50, with a siphophage morphology (Supplementary Figure S4B). Their ≈109 kb genomes (Figure 2C) are highly similar (>99%; Supplementary Figure S1A) with phage DevonBitter (Genbank: OR896848) as their closest relative. Roth39, as the representative phage of this genus, had the highest burst size of all phages in the collection, releasing 524±62 phage particles per cycle (Figure 2H). The Roth *Sugarlandvirus* also had the highest percentage of hypothetical proteins (∼72%) within the phages in the collection, proportional to their genome lengths and encoded genes (∼141/196 CDS; Figure 2C).

The Roth *Sugarlandvirus* and their relatives maintained a homologous genome architecture, except for phage Spivey (Genbank: MK630230), which has a big portion of its genome inverted, potentially due to assembly issues (Supplementary Figure S2C), and phages DevonBitter (Genbank: OR896848) and Torridgeon (Genbank: OR896843), which have a large gene deletion where a tail fibre protein is missing (Supplementary Figure S2C). This deletion may represent changes in host range for these phages.

#### Webervirus

The largest proportion (n=19/53) of the isolated KlebPhaCol phages belong to the *Webervirus* genus, *Drexleviridae* family. These siphophages (Supplementary Figure S4C) have ≈49 kb genomes and are relatively similar to each other (≥86% intergenomic similarity; Supplementary Figure S1A), with six representatives selected for further characterisation (Figure 2D). These phages show variable burst sizes ranging from 16 virions up to 200 virions released per infection cycle (Figure 2H). The Roth *Webervirus* encode ≈87 CDS of which ≈29 are annotated. These phages maintained genome synteny with their relatives, with the tail fibre protein having the least shared similarity, and that of Roth93 and PWKp14 (Genbank: MZ634345) being truncated (Supplementary Figure S2D). Most phages contained additional CDSs encoding hypothetical proteins between their two tail fibre proteins, some with no shared identity – except for Roth44, which had no additional CDSs in this region (Supplementary Figure S2D). All *Weberviruses* encoded an adenine-specific methyltransferase at the 3’ end of their genomes, but phages Roth93 and Roth85 had an additional one (of identical homology) near the 5’ end, which was absent in all other phages included in the genomic comparison. We also identified five *Webervirus* that possess a CDS (Roth09_gp33, Roth10_gp33, Roth71_gp34, Roth83_gp33, Roth84_gp33) with significant similarity (Evalue ≤1E-08) to the putative adhesin Scl1, a bacterial protein known to bind fibronectin type III domains and facilitate colonisation of the superhost (114). The DNA-modification enzyme, adenine methyltransferase (AMT) is encoded by all *Webervirus* (Supplementary Figure S3B).

#### Jiaodavirus

The second most abundant phages of KlebPhaCol (n=17/53) belong to the *Jiaodavirus* genus, *Straboviridae* family. These myophages (Supplementary Figure S4D) have large genomes of ≈167 kb and have ≥91% genome similarity between each other (Supplementary Figure S1A). Five representative phages were selected for further characterisation (Figure 2E), all of which had variable burst sizes ranging from 60 up to 379 virions released per infection cycle (Figure 2H). They have a homologous genome synteny to their relatives, but phage KM13 (Genbank: MN101229) has some deletions in the 3’ end before the ribonucleotide reductase gene product, with most of the missing genes encoding for hypothetical proteins but also a restriction endonuclease, RNA ligase, an ACL protein and a deaminase. The *Jiaodavirus* phages encoded the most DNA-modifying enzymes compared to the rest of the phages in the collection, including a glycosyltransferase family 2, two thymidylate synthases, a DNA alpha glycosyltransferase and a 5-hydroxymethyl-2’-deoxycytidine, which result in 5hmdC and 5gent-mdC DNA modifications as confirmed by HPLC (Supplementary Figures S2 and S3).

#### Slopekvirus

Seven phages of the KlebPhaCol belong to the *Slopekvirus* genus, *Straboviridae* family. These myophages (Supplementary Figure S4D) have the largest genomes in the collection (≈174 kb) and high intergenomic similarity (>99%; Supplementary Figure S1A). Their closest relative at the time of our search was phage KpnM_VAC13 (Genbank: MZ322895) (Figure 2F). Roth88 was selected as a representative *Slopekvirus* and has a burst size of 30±12 virions per infection cycle (Figure 2H). Roth88 maintained a highly homologous genome synteny with its relatives. The genes with most notable differences were tail proteins, likely with impact in the phage host range. Another notable difference was phage P-PK2 (Genbank: MT157285), with a slightly shorter genome due to a deletion of a mid-genome region of small CDSs annotated as hypothetical proteins. Interestingly, each phage may have two putative depolymerases with an unusual non-beta-helical shape, requiring further experimental validation (Supplementary Table S3). Like *Webervirus,* the *Slopekvirus* also encoded the DNA-modification enzyme AMT (Supplementary Figure S3B).

#### New *Nakavirus* genus

While most Roth phages could be classified into existing viral taxa, RothC and RothD could not be assigned to any existing viral families using standard classification tools like PhageGCN (73) and vContact2 (74). Therefore, we propose the establishment of a new order, *Felixvirales,* with a new family, *Felixviridae,* and a new genus, *Nakavirus*, to accommodate these phages. Following ICTV genome-based guidelines, we began by performing a BLASTn search against the NCBI database, but no closely related phages were found within the top 10 hits. After filtering to the virus taxa ID (10239), the closest matches were a partially complete metagenomic phage assembly: ctlJz2 (Genbank: BK029112) (98), and the previously cultured but unclassified phage vB_Kpn_Chronis (Genbank: MN013086)^76^. Due to its complete genome, only phage vB_Kpn_Chronis was selected for further analysis.

Because RothC and RothD target gut-associated ST323 *K. pneumoniae* , we expanded the search to gut phage databases. BLASTn analysis against the Gut Phage Database (GPD) (81) identified several related phages (Evalues ranging from 0 to 9E-132), but most were incomplete or of low quality. After filtering, 132 high-quality metagenomic hits were retained, which were then clustered with RothC, RothD and vB_Kpn_Chronis using vContact2, forming a coherent network that confirmed taxonomic relationship (Figure 3A).

**Figure 3.**
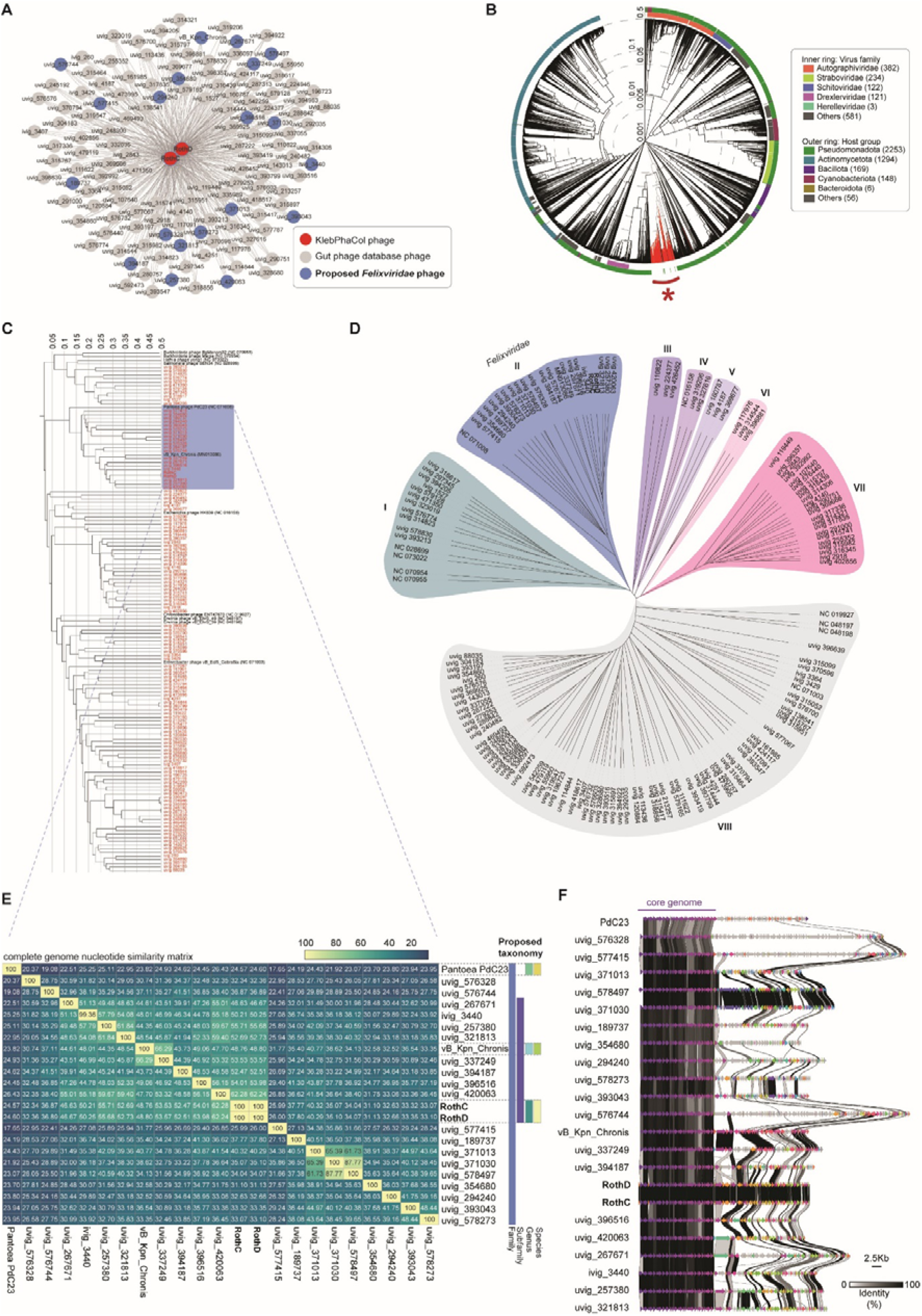
Taxonomic classification of the proposed *Felixviridae* phage family. (**A**) vContact2 network of phages RothC and RothD clustering with high confidence to phages from the Gut Phage Database. Phage vB_Kpn_Chronis was added to the network as it was identified as a previously cultured close relative of RothC and RothD. Phages that ultimately belonged to the proposed *Felixviridae* family are highlighted in blue. (**B**) VipTree circular phylogenetic tree of all phages from the cluster in a) with the dsDNA viruses from the VipTree database. The phages from the cluster in a) are highlighted in red and with the asterisk. (**C**) Pruned tree from the red cluster in b), midpoint rooted, suggesting a new monophyletic clade for the cluster and thus a new phage order, proposed as ‘*Felixvirales’*. The reference phages that fell within the cluster are coloured in black, all phages in red represent those in the cluster from a). Boxed in blue are the proposed *Felixviridae* phages. (**D**) Unrooted tree from c) highlighting the different clades within the cluster. Phages within clade VIII are highly divergent and undefined. Phages in clade II represent the proposed *Felixviridae* family. This tree was visualised using iTOL. (**E**) Whole genome nucleotide similarity matrix of *Felixviridae* phages and the proposed taxonomy below the family rank. Phage lifestyle as predicted by Bacphlip is shown below the matrix. Morphology was taken from the literature for reference phages (PdC23 and vB_Kpn_Chronis), assessed by TEM for the Roth phages, and where data was available, taken from the Gut Phage Database metadata for the remaining phages (only for uvig_576328). (**F**) Whole genome synteny plot of the *Felixviridae* phages highlighting a highly conserved core genome.

Further classification using VipTree, which relies on proteomic comparisons (82), clustered RothC, RothD, and their relatives into a distinct group (Figure 3B, red asterisk). A total of 143 phages (including RothC, RothD, and vB_Kpn_Chronis) were identified within this monophyletic clade (Figure 3C), which we propose to classify under a new order, *Felixvirales*. Based on VIRIDC intergenomic similarities, this order appears to contain seven well-defined families (Figure 3D), one of which we named *Felixviridae*, and further proposed the subfamily *Maevirinae* (demarcation criteria of ≥45%), which includes RothC, RothD, and vB_Kpn_Chronis (Figure 3C-E). At the genus level (nucleotide identity threshold of ≥70%), we suggest *Nakavirus* for RothC and RothD, and *Chronisvirus* for vB_Kpn_Chronis. Lastly, based on ≥95% nucleotide identify, we propose the species *Nakavirus sapi* for RothC and RothD, and *Chronisvirus chronis* for vB_Kpn_Chronis (Figure 3E).

The *Nakavirus* RothC and RothD have a genome size of ≈46 kb (Figure 2G) and are the only temperate phages of the collection. Interestingly, they did not match any prophages from their isolation strain 80528 or the co-cultured strain RSUH15, despite being isolated against these. Their infectious cycle was the longest of all Roth phages, reaching a plateau at 100 min (Supplementary Figure S5E), but their burst size was comparable to other Roth phages, averaging 263±32 and 116±23 particles per cycle (Figure 2H).

While RothC and RothD share little overall homology with their relatives, they have a highly conserved region from gp1 to gp24 (1-20, 241bp), encompassing structural proteins (Supplementary Figure S2G). Genomic synteny analysis of the proposed *Felixviridae* family revealed that the conserved region spanning the first 20, 241 bp is shared across all members (Figure 3F). This core region is mostly composed of structural proteins, as confirmed through alternative annotation methods like Phold(65), which provided more detailed predictions than sequence-based approaches (Supplementary Figure S8). Outside this core region, gene conservation was minimal, further emphasizing the uniqueness of these phages. Despite this uniqueness, RothC, RothD, and all but two (PdC23 and uvig_576328) of their *Felixviridae* relatives were temperate phages, with no detectable DNA-modifying genes or confirmed depolymerases. However, gp28 (annotated as tail spike receptor binding N-terminal domain) in RothC and RothD showed high scores on RBPdetect (69) and DepoScope (70), suggesting potential receptor binding functionality, but with an unusual non-beta helical shape, requiring further experimental validation (Supplementary Table S3).

### Characterisation of KlebPhaCol strains

We characterised the 74 bacterial strains in KlebPhaCol by examining their ST, KL- and O-types, focusing on prophages, virulence factors, stress resistance, antimicrobial resistance, anti-phage defence systems, and capsular locus integrity. To facilitate access to the strains and their metadata and encourage comparisons to other *Klebsiella* strains, we have deposited the *Klebsiella pneumoniae* strains into a Pathogenwatch (86) collection (see Data availability).

#### ST, KL-, and O-types

The 74 strains encompassed 40 ST-types, 32 KL-types and 10 O-antigen types (Supplementary Table S2), covering a wide range of the known diversity of *Klebsiella* in this regard. The most prevalent ST-types in KlebPhaCol include clinically relevant types associated with antimicrobial resistance, ST258 (n=8 strains), ST14 (n=8), ST11 (n=6), ST101 (n=5) and ST15 (n=4) (Figure 1E). The rest of the ST-types are covered by either one or two strains. Notably, we hold two strains of ST323, recently associated to gut colonisation and inflammatory bowel disease exacerbation (21).

Regarding KL-type, KL2 is the most prevalent KL-type in the collection (n=10 strains) and is highly clinically relevant due to its strong association with virulence traits (115, 116). The other notoriously pathogenic KL-type, KL1, is covered by two of our strains (116–118). Other common KL-types included KL24 (n=6), KL106 (n=5), and KL17 (n=5) (Figure 1E).

Lastly, KlebPhaCol strains represent ten of the 13 known O-antigens for *Klebsiella* (119) (Supplementary Table S2). Two strains also have OL102 and OL103 that are currently unclassified O-antigens. The most represented O-antigen is O1 (n=22), followed by Oafg and O2a (n=12 and 11, respectively). OL101 recently classified as a 13^th^ class of O-antigen (OL13) is found in six strains (119).

#### Prophages

Almost all strains encoded at least one prophage, except for KLEB9 with zero predicted prophages. Strain 51851 stood out with 18 predicted prophages, significantly higher than the average of 5±4 across the collection (Figure 4A).

**Figure 4.**
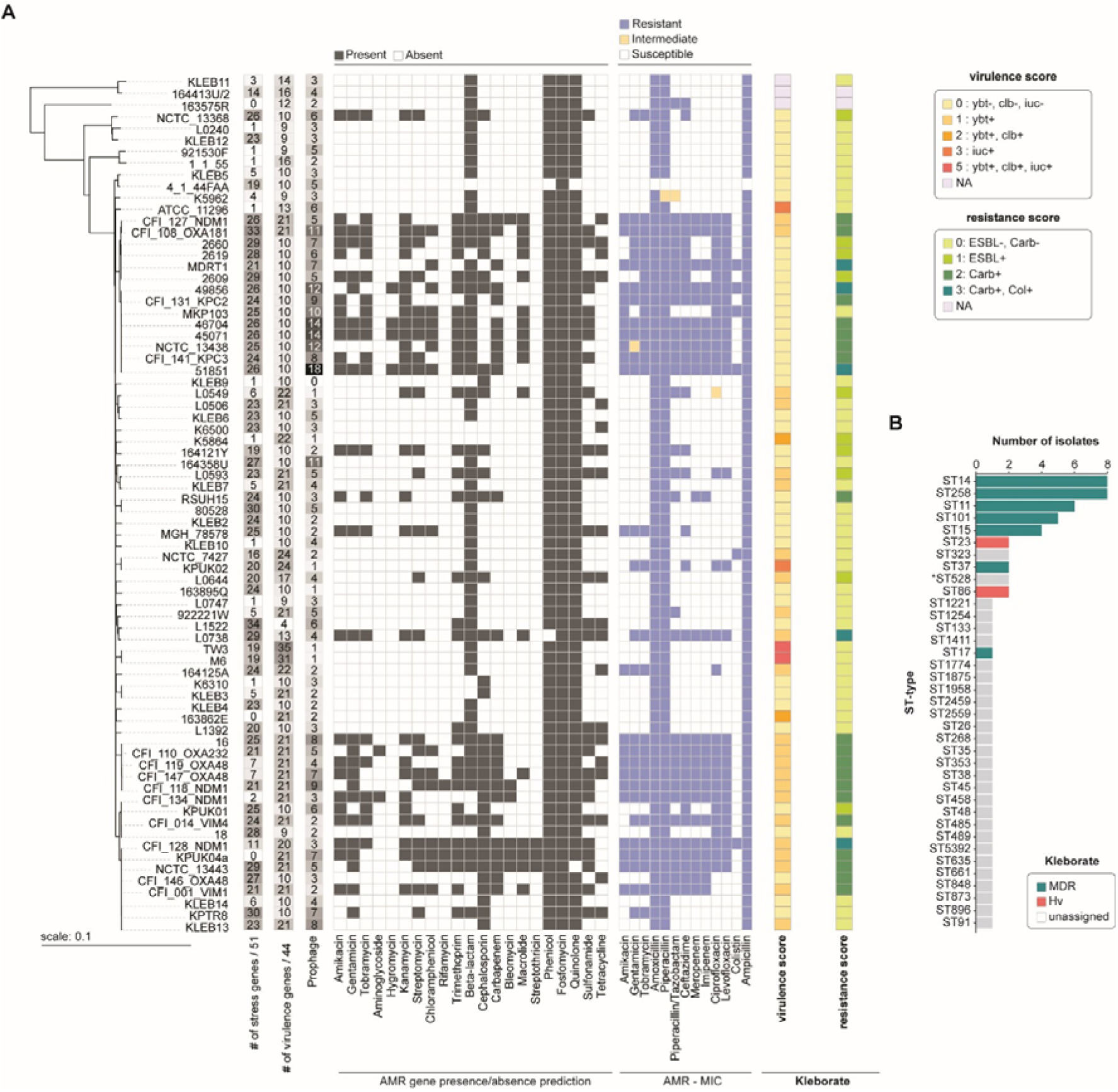
Characteristics of the 74 strains in KlebPhaCol. (**A**) Phylogeny of the 74 strains matched with the number of stress resistance genes (out of a total of 51), virulence genes (out of a total of 44), prophage predictions, and presence/absence of antimicrobial resistance (AMR) genes. The minimum inhibitory concentration (MIC) of 13 antibiotics was also tested for each strain and resistance patterns are illustrated in the coloured heatmap. Virulence and resistance scores were calculated with Kleborate. (**B**) Kleborate predictions of strains with multidrug resistance (MDR) genes and genes conferring hypervirulence are predicted by ST-type. *ST528 strain L1392 typed as ST716 by MLST typer.

#### Virulence factors

Regarding virulence, we identified 44 virulence genes, with an average of 14±6 per strain. Strains TW3 and M6 exhibited the highest predicted virulence profiles, with 35 and 31 virulence genes, respectively, earning them a Kleborate virulence score of 5 due to their encoding of all three siderophore systems (yersiniabactin, *ybt*; colibactin, *clb*; and aerobactin, *iuc*) (Figure 4A) (120). While not experimentally verified in this study, these two ST23 strains were classified as hypervirulent by Kleborate (Figure 4B) – notably, they are the two KL1 strains of the collection. Other notably predicted hypervirulent strains included ST86 strains NCTC_7427 (KL2) and KPUK02 (Figure 4B), although with lower virulence scores of 2 and 3 (Figure 4A). The most common virulence genes were *entB*, *ompA, fepC, ykgk,* and genes from the *yag* cluster (Supplementary Table S2), which contribute to enterobactin siderophore production (121), host immune evasion (122), and biofilm formation (123).

#### Stress resistance

Stress resistance genes were prevalent in the KlebPhaCol collection, with strains encoding an average of 17±6 genes. The most frequently found gene was *fief,* present in 70 out of 74 strains, responsible for iron and zinc efflux (124). While most strains had multiple stress resistance genes, strains 163575R, 163862E, and KPUK04a encoded none, and nine others contained only one (Figure 4A, Supplementary Table S2).

#### Antimicrobial resistance

Genomic analysis revealed the presence of genes potentially mediating resistance to 22 antibiotics, including six aminoglycosides, two amphenicols, and various others (Figure 4A). Most of these predictions were based on the presence of one gene (Supplementary Table S5). On average, strains had resistance genes for 8±4 antibiotics. Strain CFI_128_NDM1 carried resistance genes for 18 antibiotics, the highest in the collection, while strain 4_1_44FAA carried the least resistance genes. Accordingly, the Kleborate resistance score for CFI_128_NDM1, as well as to L0738, 51851, 49856 and MDRT1, was 3 (highest), indicating likely genotypic resistance to carbapenems and colistin, both last-line antibiotics (Figure 4A). High carriage of genes associated with resistance were observed for phenicols, quinolones, beta-lactams (in general, including cephalosporins), and trimethoprim (Figure 4A). Contrastingly, streptothricin, rifamycin, hygromycin and the aminoglycosides emerged as the antibiotics for which there were the fewest predicted resistance genes.

These analyses do not necessarily predict phenotypic resistance, with the possibility of resistance being mediated by genes operating in a multifactorial manner and intrinsic resistance associated with poor cell penetration and/or efflux. Therefore, experimental validation of these predictions was carried out for a defined selection of clinically important antibiotics using minimal inhibitory concentrations (MIC). These confirmed that strain 4_1_44FAA had the lowest resistance, while CFI_128_NDM1, MDRT1, 49856, and 51851 were resistant to all tested antibiotics, consistent with their high Kleborate resistance scores. Carbapenem resistance predictions were 100% accurate, but resistance to gentamicin and tobramycin was higher in laboratory conditions than predicted (23 vs 28 strains and 22 vs 31 strains, respectively), and amikacin resistance was slightly lower than anticipated (23 vs 25 strains, Figure 4A and Supplementary Table S2). This demonstrates the difference between genotypic resistance predictions and phenotypic susceptibility determination.

#### Defence systems

The strains in this collection encode a total of 93 distinct defence systems, with an average of 11±4 systems per strain (Supplementary Figure S9A). Most systems were rare, with 54 out of 93 systems present in fewer than five strains. Only RM type IV and AbiE systems were found in approximately 85% of the strains. Other notable defence systems included Mok Hok Sok, RM types I and II, and SoFic, present in ≥50% of the strains (Supplementary Figure S9A). Defence system distribution varied per strain, with certain systems associated with specific ST-types. For example, the AbiU system is primarily found in ST11 and ST258 strains, while DISARM, DRT2, Kiwa, Mokosh type I, and PifA are almost exclusively found in ST23 strains (Supplementary Figure S9A, Supplementary Table S2). Dpd is only present in ST323 strains, while GAO_19 and PD-T7-2 are also almost exclusive to these. Strain L0738 has the highest number of defence systems (n=31), while KLEB11 has only two (Supplementary Figure S9A).

### Roth phages infect up to 19 KL-types but host range varies with bacterial growth medium

Phage host range in *Klebsiella* is largely dictated by the presence or absence of the capsular polysaccharide (33, 34), which can be influenced by media composition (125–127), and as a result affect phage infectivity. To assess possible influences of media in phage host range, we performed host range assays in tryptic soy broth (TSB) and lysogeny broth (LB), both commonly used in *Klebsiella* research and with different nutritional compositions (33, 45, 128–130).

The Roth phages demonstrated a broad ability to infect a wide range of *Klebsiella* strains, with notable success across multiple sequence types (ST), capsular types (KL), and O-antigen types (Supplementary Figure S10). Across the 74 strains tested, 46 (62%) were susceptible to the Roth phages (Figure 5A), including 27/40 (67%) ST-types, 28/32 (88%) KL-types, and 9/13 (69%) O-antigen types. As expected, susceptibility was highest among the strains used for phage enrichment during isolation, which includes the clinically-relevant strains ST258, ST14, and ST11—associated with varying KL-types and typically O-antigens O1 and O2afg (Supplementary Figures S6 and S7).

**Figure 5.**
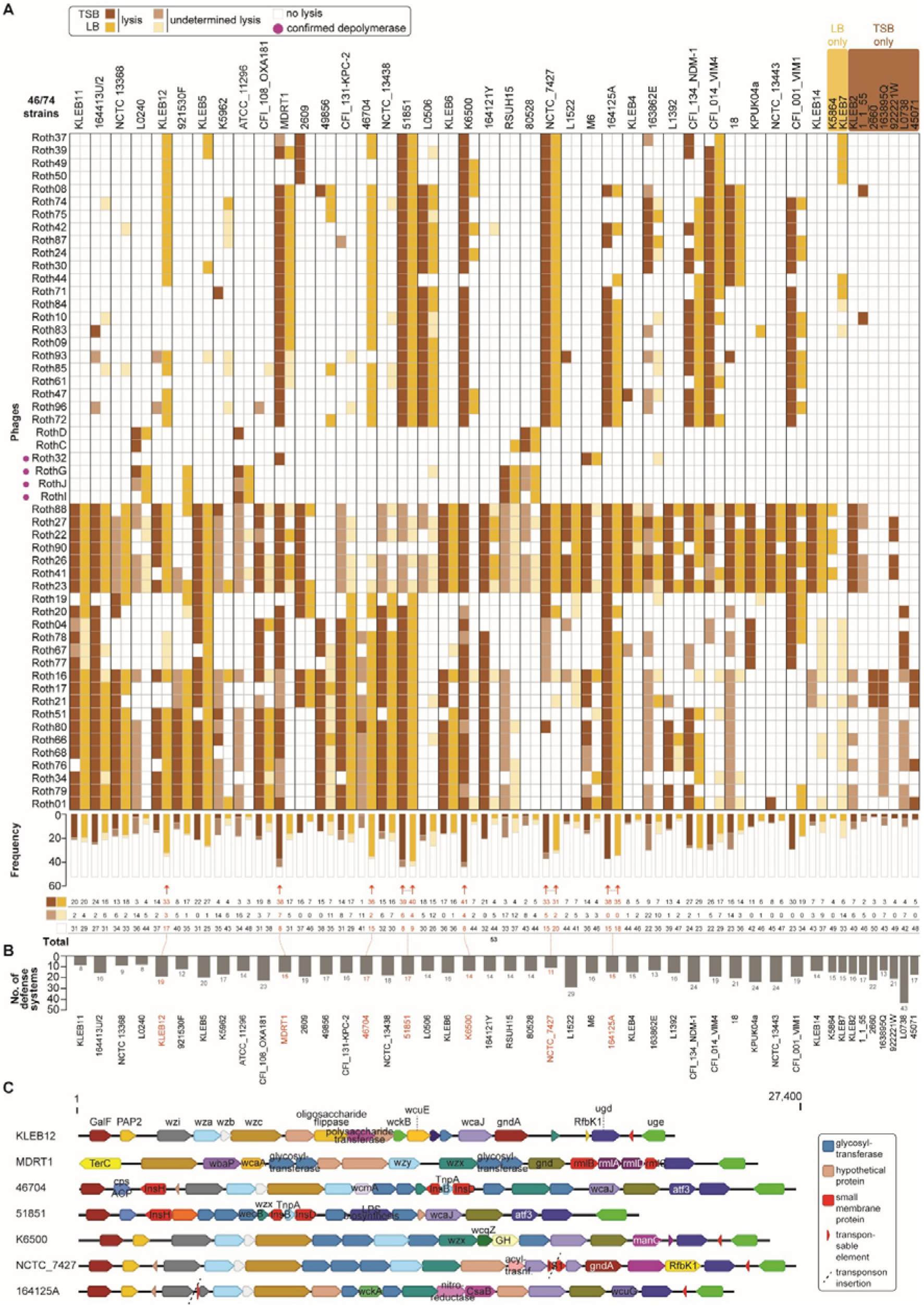
Host range of the 53 Roth phages. (**A**) Heatmap illustrates the infection patterns of the 53 phages against the susceptible strains (n=46 out of 74 tested), in both TSB (brown) and LB (yellow) media. Undetermined lysis is marked by opaque squares while no lysis is represented by white squares. All infection patterns (lysis, undetermined lysis, no lysis) are quantified below the heatmap and highly susceptible strains are illustrated in red and highlighted with a red arrow. Phages are ordered phylogenetically. (**B**) Total counts of defence systems harboured by the susceptible strains. Those strains with high susceptibility highlighted in a) are also marked in red here. (**C**) Assembled capsule loci of the strains marked in red in a) and b). Gene deletions are noted by the empty genes and transposon insertions are highlighted with a red aura.

#### *Slopekvirus* have the broadest host range

Phages from the *Slopekvirus* genus were the most effective, with Roth23, Roth41, Roth26, Roth22, and Roth88 infecting 18-20/74 (∼26%) strains in LB (Supplementary Figure S6), and 24-25/74 (∼33%) in TSB (Supplementary Figure S7). The *Jiaodavirus* Roth01 also exhibited a broad host range, infecting 17/74 (23%) strains in LB and 21/74 (28%) in TSB.

In terms of ST-type infectivity, *Slopekvirus* were able to target between 11-14 ST-types in LB and 15-16 in TSB (Supplementary Figure S10A, B). Interestingly, ST258 strains are resistant to *Slopekvirus*, but efficiently targeted by all *Jiaodavirus*, *Sugarlandvirus*, and *Webervirus* .

*Slopekvirus* also displayed the broadest range in terms of KL-types. Phages from this genus infect 12-16 KL-types in LB and 17-19 in TSB (Supplementary Figure S10C, E). *Slopekvirus* exhibited significant success against KL2 strains, infecting a large proportion of these clinically relevant types.

Roth phages were able to infect strains associated with nine of the ten known O-antigen types in the KlebPhaCol collection, except for O12 (Supplementary Figure S10D, F). Phages were most successful against strains with O-antigen O2afg, although *Slopekvirus* were more efficient at targeting O1, O2a, and O3/O3a strains, while *Jiaodaviruses* excelled at targeting O2afg. In LB, Roth23 and Roth88 in particular, infected strains associated with 8/10 known O-antigen types, in addition to unknown OL-type OL103 - further confirming the broad host range of these *Slopekvirus*.

#### Phage infectivity is influenced by bacterial growth media

The impact of media composition on phage infectivity was evident in the differential success rates observed between TSB and LB (Figure 5A). In TSB, 645 total phage infections were recorded, compared to 541 in LB, indicating higher infectivity overall in TSB (Supplementary Figures S6 and S7). The media-specific differences were particularly pronounced for certain strains. For instance, strains KLEB12 and 46704 had more than double the amount of targeting phages in LB, while MDRT1 and K6500 were 2x and 5x more susceptible in TSB, respectively (Figure 5A). Moreover, some strains were only infected in one of the two media. For example, K5864 and KLEB7 were only infected in LB by four and six phages, respectively, while strains KLEB2, 1_1_55, 2660, 163895Q, 922221W, L0738, and 45071 were exclusively infected in TSB (Figure 5A).

Interestingly, phage infectivity varied by genus in response to media composition. *Jiaodavirus*, *Slopekvirus*, *Nakavirus*, and *Webervirus* showed higher success rates in TSB, with ∼1.3x more infections recorded in this medium. For example, *Slopekvirus* Roth26 and Roth27 can infect up to 19 different KL-types in TSB, whereas in LB they can only infect 14 and 13 different KL-types, respectively (Supplementary Figure S10C, E). In contrast, capsule-targeting phages such as Roth32, RothG, RothJ, and RothI were twice as effective in LB compared to TSB (Supplementary Figures S6, S7, and S10). This discrepancy could be attributed to enhanced capsule production in LB, a nutrient-rich medium (127), which for capsule-targeting phages, would have improved phage adsorption by making the capsule more accessible. In contrast, capsule production is likely reduced in the more carbon-limited TSB medium (due to it containing soytone instead of yeast extract) (127), potentially uncovering other surface receptors that phages may instead rely on such as the O-antigen, FhuA, or other membrane components.

While *K. pneumoniae* is the most pathogenic species among *Klebsiella* spp., other species are emerging with serious pathogenic concerns (131, 132). KlebPhaCol phages demonstrated lytic activity beyond *K. pneumoniae* , including *K. oxytoca* (164413U/2, KLEB11), *K. similipneumoniae* (NCTC_13368), *K. variicola* (921530F and 1_1_55), and *K. pneumoniae sub ozaenae* (ATCC_11296) (Figure 5A). Together this indicates that the KlebPhaCol collection could be expanded to study interactions across a broader range of *Klebsiella* species.

#### Abundance of defence systems does not correlate with phage susceptibility

Antiphage defence systems pose a barrier to phages once inside the cell (133–135). We observed that some of the least susceptible strains like L1522 and L0738, harboured a disproportionately high number of encoding defence systems when compared to the rest of the strains (22 and 31, respectively; Figure5B). To investigate whether phage susceptibility was associated with the number of defence systems in each strain, we calculated and visualised Spearman rank correlations between the number of defence systems and various infection outcomes, including productive infections, no infections, and undetermined infections (i.e., ‘lysis from without’) (Supplementary Figure S9B). Our analysis revealed that in the panel of phage and clinical isolates tested, there is no significant correlation between the number of encoded defence systems and phage susceptibility, regardless of the media type. The recently reported PhageHostLearn model for *K. pneumoniae* phages suggests that RBP variability accounts for most of the host spectrum diversity (136), hinting at a lesser role of phage defence systems in shaping host range. However, conflicting findings have been reported for non-capsulated species (137), and further investigation is needed. Specifically, future analysis of phage adsorption to strains without productive infection may provide new insights and uncover correlations not evident with the current dataset.

Additionally, we predicted the putative anti-defence proteins harboured by the Roth phages to explore whether these might influence phage infectivity. We identified three putative anti-defence genes in the *Jiaodavirus* and *Slopekvirus* phages (Supplementary Table S1). In *Jiaodavirus*, these included two anti-CBASS and one anti-TA, while the *Slopekvirus* encoded two anti-CBASS and one anti-RM. We also identified an anti-RM gene in a subset of *Webervirus* phages, specifically those branching from the second clade within the *Webervirus* group (Figure 1B). Interestingly, phages encoding anti-defence genes were not notably more effective in infecting strains compared to other genera members (Supplementary Figures S6 and S7).

#### Capsule alterations contribute to a broader phage host range

For most reported *Klebsiella* phages, the capsule is the primary surface receptor to which they attach (32, 34, 40, 45), although other surface receptors like the O-antigen and the LPS have now also been shown to serve as primary receptors for some *Klebsiella* phages (33, 34, 140). We observed that some highly phage-susceptible strains (MDRT1, 51851, 46704, NCTC_7427 and 164125A) contained disruptions that may affect capsule production (Figure 5C, Supplementary Figure S11). For example, strains MDRT1 and 51851 showed large deletions at the 5’ end of the capsule locus, affecting genes such as *wzi*, *wza*, and *wzb* (and *wzc* in 51851), which are essential for capsule polymerisation, extension, and export (141) (Supplementary Figure S11B, D). Strain 46704 contained several transposable elements in its capsule locus, while strains CFI_014_VIM4, KPUK04a, 164125A, NCTC_7427, and KLEB5 had transposon insertions in the *wcaJ/wbaP* or *wzi* genes, which could disrupt capsule production and export. Indeed, Kaptive v3 reports ‘Capsule null’ for several of these stains and others used during phage isolation (Supplementary Table S2). However, other highly susceptible strains, such as K6500 and KLEB12, had no capsule disruptions, suggesting that other factors may account for their high susceptibility.

Some of the strains with disrupted capsule loci were used as isolation hosts for 19 of the 53 KlebPhaCol phages (Supplementary Figure S11, Supplementary Table S1). This led us to hypothesise that these phages do not rely on the capsule as their primary receptor, which may contribute to their broader host range. In contrast, phages Roth32 (*Gajwadongvirus*), RothG, RothI, and RothJ (*Drulisvirus*) exhibited narrower host range and were isolated on strains with intact capsule loci (M6 and RSUH15; Supplementary Figures S6 and S11E, F). Together with evidence that these phages encode depolymerases, this suggests they likely target the capsule as their primary receptor.

For phage families infecting strains with disrupted capsule loci, we further analysed bacteriophage insensitive mutants (BIMs) that emerged after phage exposure on the capsule-deficient strain 51851. Culturing and sequencing these BIMs, followed by re-testing phages infectivity (Supplementary Figure S12), revealed that KlebPhaCol phages from the *Sugarlanvirus*, *Webervirus*, *Slopekvirus*, and *Jiaodavirus* families use either the O-antigen of lipopolysaccharide, outer membrane protein FhuA, or both as receptors (Supplementary Figure S12). These findings confirm that these phages are capsule-independent.

### Roth phages target IBD-related ST323 K. *pneumoniae* in aerobic and anaerobic conditions

Certain *K. pneumoniae* ST-types appear linked to specific infection contexts, such as ST323, which has been implicated in inflammatory bowel disease (IBD) (21). Federici and colleagues demonstrated that IBD patients who had enriched populations of *K. pneumoniae* in their gut also experienced exacerbated inflammation and worse symptoms. Notably, the administration of phages ameliorated the inflammation and population of *K. pneumoniae* in a mouse model. Given the relevance of *K. pneumoniae* as a gut pathobiont, we aimed to isolate phages effective against ST323, a now confirmed key clonal group in the human gut. Five phages in KlebPhaCol – *Nakavirus* RothC and RothD, and *Drulisvirus* RothG, RothJ and RothI – were isolated as a result. These phages share low genomic similarity with those used by Federici and colleagues (21) (< 39%) and exhibit narrow host ranges, infecting the two ST323 (KL21) strains (RSUH15 and 80528) as well as three non-ST323 strains (ST91, ST635, and ST1875) (Figure 6A).

**Figure 6.**
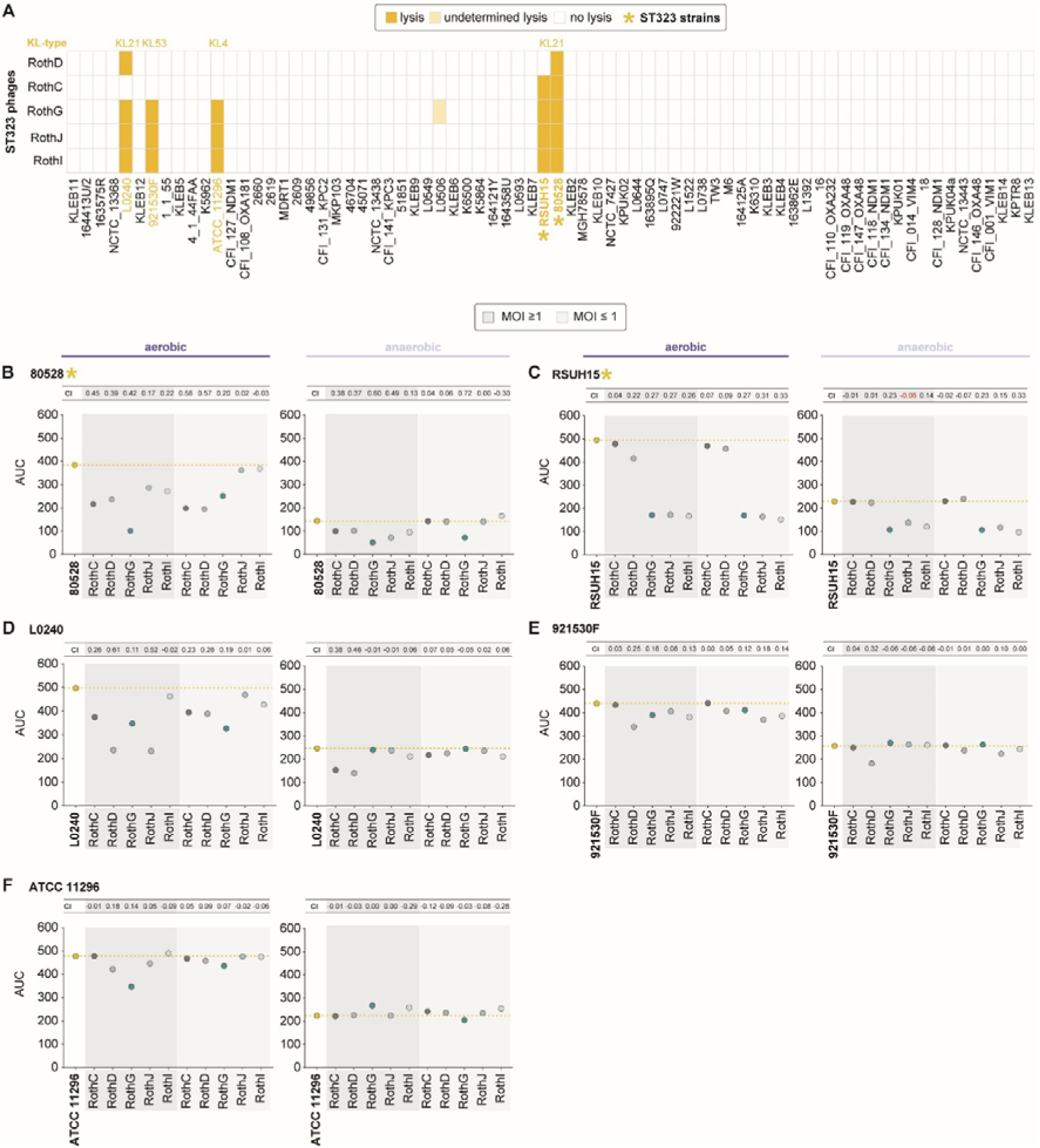
Infectivity of ST323-targeting Roth phages in aerobic and anaerobic conditions. (**A**) Heatmap of the infectivity patterns of the ST323-targeting Roth phages in LB medium. The capsule locus (KL) type of susceptible strains is written above, and ST323 strains are marked with an asterisk. (**B-F**) Growth curves of the ST323-targeting phages against the susceptible strains in aerobic and anaerobic conditions, tested using two multiplicities of infection (MOI, >1 and <1), and represented as area under the curve (AUC) plots with centroid index (CI) values displayed above. A CI value closer to 1 indicates better phage efficiency. Bacteria controls are marked in yellow.

Given the facultative anaerobe nature of *K. pneumoniae* and its adaptation to the gut, we tested whether these phages could retain activity in both aerobic and anaerobic environments. Using liquid infection assays under both conditions, at two multiplicities of infection (MOI), we evaluated phage efficacy against ST323 strains (Supplementary Figure S13, Supplementary Table S6). Results were represented using both the area under the curve (AUC) and centroid index (CI) values (48) to enhance interpretability (Figure 6B-F). CI values are a metric that accounts for fluctuations overtime in phage killing liquid assays attributed to phage-host dynamics, by considering the distribution of the entire bacterial growth curve, specific to phage-bacterium dynamics (48). A CI value of 0 indicates the phage behaved as the control, a negative CI value indicates the bacterium grew past the control, while a value closer to 1 indicates a successful bacterial growth inhibition persistent overtime (48).

As expected, bacterial growth was limited under anaerobic conditions, but several Roth phages remained active (Figure 6B, C). Notably, RothG, RothJ, and RothI effectively targeted RSUH15, albeit with reduced efficiency compared to aerobic conditions. For strain 80528, all phages displayed activity under anaerobic conditions at high MOI, with RothG being particularly efficient even at lower MOI. Amongst the non-ST323 strains (Figure 6D, E, F), RothC and RothD were the most effective in anaerobic conditions, especially RothD, which successfully infected strains L0240 and 921530F when used at higher MOI. However, none of the phages efficiently targeted strain ATCC_1129 in anaerobic conditions, likely due to their already reduced activity under aerobic conditions. The CI values were strongly associated to the AUC observations. Overall, these findings suggest that the ST323-targeting Roth phages hold potential for therapeutic use in the anaerobic environment of the human gut, potentially providing a new tool for treating IBD-associated *K. pneumoniae* infections.

### *Felixvirales* is a gut-associated phage order

#### Nakavirus are associated with Enterobacteriaceae

Given that most *Felixviridae* phages have a predicted temperate lifestyle we wanted to assess their prevalence in bacterial genomes. We analysed 64, 364 complete bacterial genomes from the Bacterial and Viral Bioinformatics Resource Center (BV-BRC) database for homologues of RothC and RothD and found that all matches (n=7, 605 total; n=708 significant; Supplementary Table S7) were exclusive to the *Enterobacteriaceae* family.

*Klebsiella* species were the most common hosts (566/708; 80%), spanning eight species and 111 ST-types, with ST231 appearing most frequently (127 hits). The second most represented species was *Salmonella enterica* (66 hits, 9%). These phage-host associations were found across 67 countries, indicating a widespread global presence of these prophages. Host metadata reveals that most isolates (521) were derived from humans, though samples from hosts like chickens (10), pigs (7), termites (4), sheep (3), cattle (2), and even hedgehogs, sealions and birds suggest a broader superhost range (Supplementary Table S7). However, the location of the isolate within these other organisms remains unknown.

We also examined predicted prophages within these genomes to determine whether our identified hits were located within these regions. Of these hits, only two were entirely within prophage regions. The majority were either outside prophage regions (n=431/708; 61%) or partially within one (n=266/708; 36%). To further investigate the gene neighbourhood of hits outside prophage regions, we extracted the ten upstream and downstream genes from each hit. In 259/431 (60%) of the cases, a lysozyme gene was found within 10 CDS upstream of the hit region. Additionally, in seven of these 259 cases, an integrase gene was also identified within 10 CDS downstream of the hit region. This suggests that some hits may indeed reside within prophage regions that are not detected by the prophage identifier tool, highlighting the importance of examining gene neighbourhoods for more comprehensive analysis.

#### *Felixvirales* are gut-residing phages

The taxonomic characterisation of the *Felixvirales* RothC and RothD described above, suggests that *Felixvirales* are present in the mammalian gut. Several gut-related phage families and even orders have now been established, including the *Flandersviridae* and *Quimbyviridae* (142), Gubaphages (81), *Grandevirales* (143), and *Crassvirales* (144). The latter is the most abundant phage order within the mammalian gut known to date (144, 145), and is known to persist overtime potentially via several acquired evolutionary advantages (146, 147). To determine the predominance of *Felixvirales* in the human gut, we looked at their prevalence in the Gut Phage Database (GPD). We found a total of 1, 298 unique hits across 66 isolates and 873 metagenomes, corresponding to 0.9% of the phage genomes in the GPD (Figure 7A; Supplementary Table S8). These phages appear globally widespread, with metagenomic samples collected from 24 different countries (Figure 7A). Consistent with the analysis of the Bacterial and Viral Bioinformatics Resource Center (BV-BRC) database above, analysis of GPD metadata confirmed that these phages are restricted to hosts of the *Enterobacteriaceae* family, with *Klebsiella* spp. being the most common (510/539 hits with available host-predicted data).

**Figure 7.**
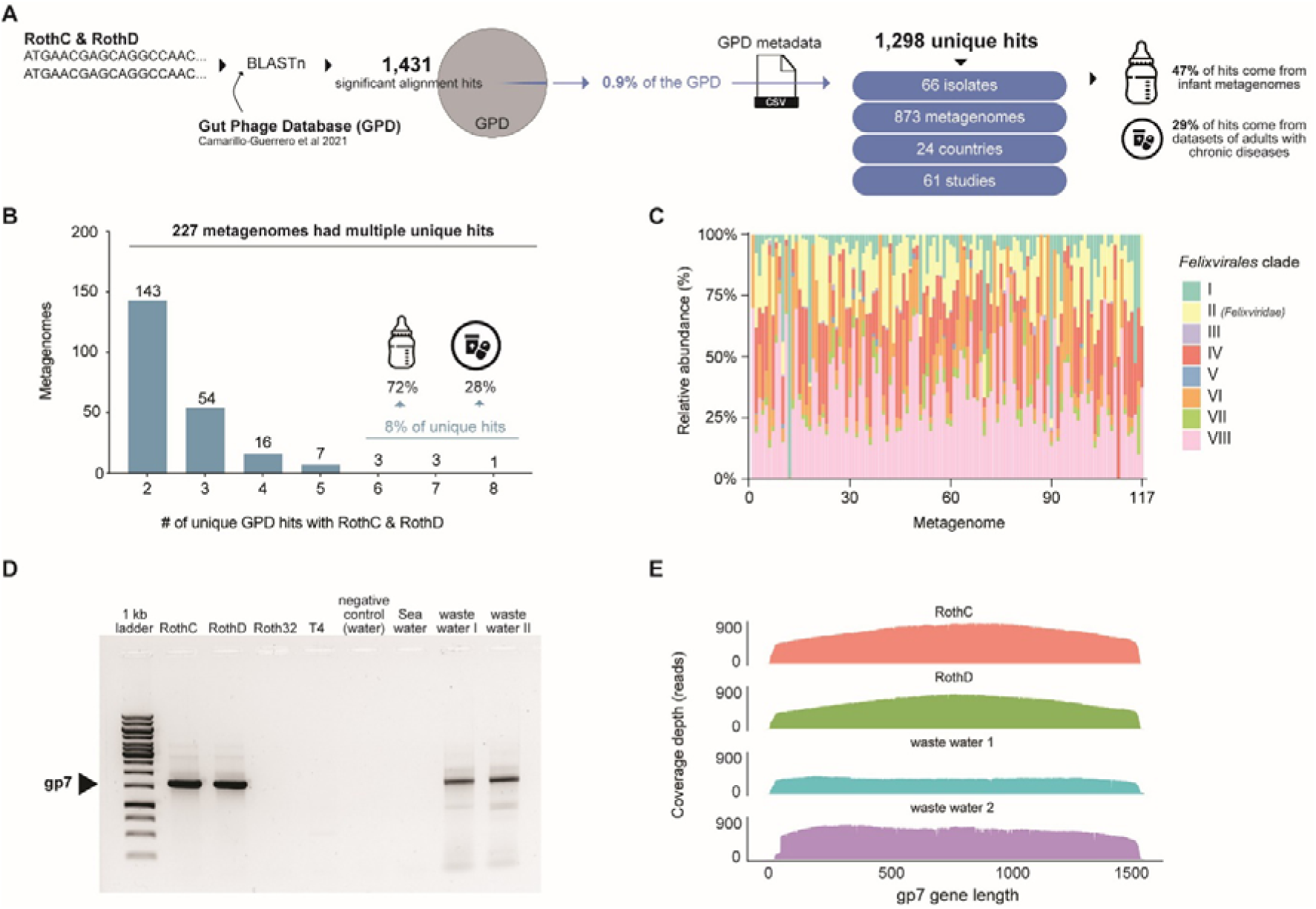
Analyses of *Felixvirales* prevalence. (**A**) Gut Phage Database (GPD) hits to RothC and RothD and their associated data taken from the GPD available metadata. (**B**) Bar plot quantifying the metagenomes containing multiple unique viral contig hits to RothC and D and the cohort type associated to some of them. (**C**) Relative abundance distribution of *Felixvirales* in the 117 metagenomes from healthy adults. (**D**) Primers to the gp7 (putative virion structural protein) of the core region of RothC and RothD can detect related phages in environmental samples by PCR. (**E**) Positive PCR products from e) sequenced by long-read sequencing and displayed as coverage depth when mapped to the gp7.

*Felixvirales*-related sequences were predominant in infant and adult cohorts, representing 92% of hits (Supplementary Table S9) and spanned both healthy and disease-associated microbiomes, highlighting their prevalence across ages and health statuses. In several (227/873) metagenomes, multiple *Felixvirales*-related phages were present, suggesting that individuals could harbour diverse populations of these phages. (Figure 7B). For example, one infant gut sample contained eight unique hits, underscoring the extensive variability of *Felixvirales* within individual gut microbiomes.

When examining disease associations, *Felixvirales* appeared in cohorts(98) with chronic conditions like obesity, IBD, and rheumatoid arthritis (Supplementary Table S10). However, no significant associations were found, suggesting these phages persist in various gut environments without clear links to disease states.

#### *Felixvirales* reside in free phage and prophage formin the gut

To assess the relative abundance of *Felixvirales* in the human gut, we analysed 117 healthy stool samples from the Human Microbiome Project (148). This analysis detected *Felixvirales* sequences in 100% of samples, with clades IV, II (*Felixviridae*), and VI being most prevalent (mean percentage of reads aligned: 0.0017%, 0.00013%, and 0.00008%, respectively) (Figure 7C, Supplementary Figure S14). These phages constituted a minor but consistent fraction of the gut microbiome. After removing bacterial reads, *Felixvirales* remained detectable in only 49% of samples, indicating many *Felixvirales* reside as prophages in bacterial cells in the gut.

#### *Felixviridae* are detectable in the environment

To assess the feasibility of PCR detection of *Felixviridae* in environmental samples, we designed primers against the conserved putative virion structural protein gp7 of the *Felixviridae* family. We screened for these phages in environmental samples collected from a wastewater facility in Portswood and Petersfield, and from the sea of the Isle of Wight (all UK-based). *Felixviridae* were present in the two sampled wastewater facilities but not in the sea water sample (Figure 7D), which we confirmed by long-read sequencing of the PCR products (Figure 7E). These results support the link of *Felixviridae* to human-associated environments and prove the feasibility of their detection in environmental samples by PCR.

Together, these data suggest *Felixvirales*, particularly *Nakavirus*, are widely distributed across diverse human and animal hosts, persisting throughout life stages and health conditions. This broad host range, coupled with environmental presence, positions *Felixvirales* as integral yet understudied components of the gut virome, meriting further exploration for their ecological and potential clinical impacts.

## Discussion

Here we introduce KlebPhaCol, a fully characterised, open-source collection containing 53 phages targeting *Klebsiella* spp., covering seven genera, and 74 *Klebsiella* strains spanning 40 ST-types and 32 K-types. By offering a centralised, no-cost, and well-documented collection, KlebPhaCol democratises access to essential resources for researchers worldwide.

Characterisation of KlebPhaCol phages reveals that Roth phages effectively target 28 known KL-types and 27 ST-types, including some of the most clinically significant strains, such as those of ST258 (carbapenamase producers (149)) and KL2 (hypervirulent (117)) types. Additionally, KlebPhaCol phages demonstrate effectiveness against other prevalent pathogenic strains, including ST11, ST23, ST14 and ST15 (12, 19, 150), and species beyond *K. pneumoniae* that are emerging as public health concerns (131, 132). KlebPhaCol also includes phages highly specific to gut-colonising *K. pneumoniae* ST323, which has been associated to inflammatory bowel disease exacerbation (21). Thus, KlebPhaCol phages can be applied to clinically relevant investigations concerning phage therapy and beyond.

Our findings further underscore the diversity of *Klebsiella* phage tropism, with KlebPhaCol containing both capsule-dependent and non-capsule-targeting phages. While capsule-targeting phages are useful in applications where precise targeting is required, we observed that non-capsule targeting phages tend to have broader host ranges, an insight supported by recent studies (33, 34, 125). This has significant implications for phage therapy: broad-spectrum phages could offer versatile treatment options, especially when capsule types vary widely among infections. Furthermore, we found that phage infectivity patterns vary by media type, likely due to environmental-driven changes in capsule production (126, 127). These results suggest that tailoring phage therapy to specific infection environments – taking into account nutrient availability and other physiological conditions at the infection site – could improve phage efficacy.

With KlebPhaCol, we identified a novel gut-associated phage order, with the proposed name of *Felixvirales*. This order, represented by KlebPhaCol phages RothC and RothD, appears to include at least seven families, with *Felixviridae* phages being geographically widespread and present across human age groups, from pre-term infants to adults. *Felixviridae* have a notable association with *Enterobacteriaceae* members, which include key human pathogens. While we did not find direct associations with disease states, the presence of these phages in healthy gut microbiomes highlights their role in the human gut virome and points to new opportunities to study gut microbiome dynamics. Additionally, other *Klebsiella*-targeting phage groups, such as *Webervirus*, have been identified as gut-associated phages that infect *Klebsiella* species (151). These findings underscore the importance of exploring the roles of various *Klebsiella* phage groups within the gut microbiome, with KlebPhaCol’s inclusion of *Felixviridae* and *Webervirus* helping facilitate further research on their prevalence and functions. By establishing PCR detection protocols for *Felixviridae*, our findings lay the groundwork for future research on these phages, both in clinical and environmental contexts.

Altogether, KlebPhaCol provides a comprehensive, openly accessible resource for studying *Klebsiella* phage interactions. It offers a uniquely broad scope, spanning critical pathogenic strains, non-capsule-targeting phages with versatile applications, and a newly described gut phage order with potential implications for gut health. Recognising the lack of standardised open biosharing regulations and pipelines, we actively participate in discussions to address this crucial need for progressive research (152). We hope that KlebPhaCol will not only facilitate new discoveries in microbiology and therapeutic research but also inspire contributions from the broader scientific community to further expand and improve this evolving resource.

## Data availability

Phage assemblies from this study are available at GenBank accessions 657785-657835, PP934563 and PP934564. Phage raw reads are available at PRJNA1192413, and annotation files at Figshare (DOI: https://doi.org/10.6084/m9.figshare.27794211.v1). Bacteria assemblies can be found under BioProject accessions PRJNA1123654, PRJNA73191, PRJNA1121092, PRJNA1121093, PRJNA31, PRJNA745534, and PRJNA1187231. Bacteria raw reads and annotation files are available at Figshare (DOI: https://doi.org/10.6084/m9.figshare.27759627.v1). Network files from Supplementary Figure S1B and Figure 3A can be accessed via NDEx at https://www.ndexbio.org/viewer/networks/5b968101-45e2-11ef-a7fd-005056ae23aa and https://www.ndexbio.org/viewer/networks/d66e5703-6463-11ef-a7fd-005056ae23aa, respectively. The KlebPhaCol strain collection in Pathogenwatch can be accessed at https://pathogen.watch/collection/tdtgvdxs4xhw-klebphacol-kp-strains. All other supporting data of this study are available within the article or through the supplementary data files. Bacterial strains and phages from this study are freely available upon request through our website, www.klebphacol.org. Please note that shipping costs and any additional requirements will apply.

## Supplementary data

Supplementary data are available at NAR Online.

## Author contributions

Conceptualisation, F.L.N.; Methodology – strain isolation, M.W., M.S., P.J.H, A.M., E.S., F.D., K.S., S.F.; Methodology – phage isolation and imaging, one-step growth curves, D.R.R.; Methodology – phage host-range, D.R.R., M.A., M.H., A.R.C.; Methodology – DNA extractions, D.R.R., C.K., F.M., Y.W., M.W.; Methodology – phage genome assembly and annotation, D.R.R., K.L., P.W., Y.L; Methodology – experimental verification of DNA modifying enzymes, P.W., Y.L; Methodology – RBP and depolymerase predictions, A.L., I.D.A., D.B., Y.B.; Methodology – strain and capsule locus assembly, M.W.; Methodology – 51851 BIM data, M.W.; Methodology – strain genome analyses, S.K.K., K.L., D.R.R.; Methodology – BV-BRC and GPD database analyses, K.L., D.R.R; Methodology – strain antibiotic susceptibility tests, P.J.H., M.W.; Data analysis and curation, D.R.R.; Writing – Original Draft, D.R.R, F.L.N.; Writing – Review & Editing, all authors; Supervision, F.L.N.; Funding Acquisition, F.L.N., J.M.S., S.J.J.B., Y.B.

## Acknowledgements

Strains used for phage isolation were provided by UKHSA, acknowledging Dr Katie Hopkins of the AMR Reference Laboratory at Colindale and we gratefully acknowledge Dr Daniela Centron for provision of ST258 *K. pneumoniae* strains from Laboratorio de Investigaciones en Mecanismos de Resistencia a Antibióticos IMPAM (UBA/CONICET), Facultad de Medicina, Universidad de Buenos Aires, Argentina. We thank Saul Faust from the University Hospital Southampton, for providing strains to the collection. We thank Qiagen for providing a 1-year license of CLC Premium Workbench to DRR. We thank Dorah Mbang and Florence Rushton from Peter Symonds College, Thomas C. Todeschini and Robyn Pringle for their laboratory assistance with some of the assays. We thank Matt Irwin, Rachel Buchanan and Sarah Aveline for the processing of the raw effluents collected in Southampton, UK. We gratefully acknowledge IRIDIS infrastructure for high-performance computing and the Biomedical Imaging Unit for the TEM facilities and technical support.

