## Supplementary Figure for "KlebPhaCol: A community-driven resource for Klebsiella research identified a novel gut phage order associated with the human gut"

### Table of Contents

**Supplementary Figure S1.** Comparisons of the KlebPhaCol phages.

**Supplementary Figure S2.** Genome synteny plots of representative Roth phages with relatives.

**Supplementary Figure S3.** DNA-modifying enzymes of Roth phages.

**Supplementary Figure S4.** Transmission electron microscope images of all KlebPhaCol Roth phages.

**Supplementary Figure S5.** One-step growth curves and burst size of all representative phages.

**Supplementary Figure S6.** Complete phage host range in LB medium.

**Supplementary Figure S7.** Complete phage host range in TSB medium.

**Supplementary Figure S8.** Core genome annotations of RothC and RothD.

**Supplementary Figure S9.** Bacteria-encoded defence systems and correlation analyses with infectivity patterns.

**Supplementary Figure S10.** Roth phage host range in terms of ST- KL- and O-antigen types.

**Supplementary Figure S11.** Assembled capsule loci of isolation and production host strains.

**Supplementary Figure S12.** Bacteriophage insensitive mutants (BIMs) reveal the receptors of selected phages.

**Supplementary Figure S13.** Growth curves of ST323-targeting phages in susceptible strains.

**Supplementary Figure S14.** Relative abundance of *Felixvirales.*

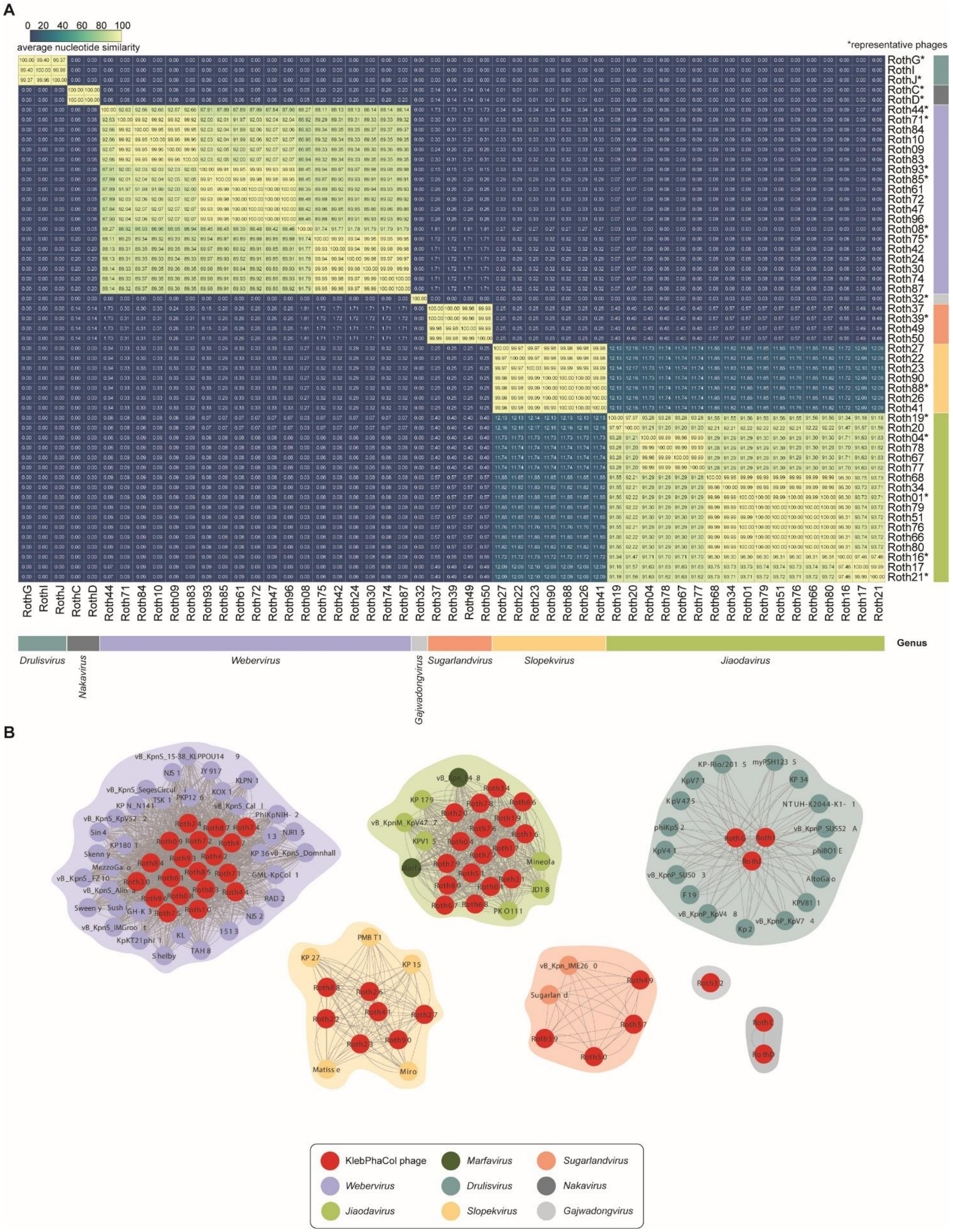


**Supplementary Figure S1.** Comparisons of the KlebPhaCol phages. (**A**) Intergenomic similarity matrix of the Roth phages calculated using VIRIDIC on the web server and visualised in Rstudio using pheatmap. Genera are grouped by colour and representative phages are marked with an asterisk. (**B**) vContact2 network of the Roth phages (red nodes) with the *Klebsiella* phages of the vContact2 database. Phages are clustered by genus.

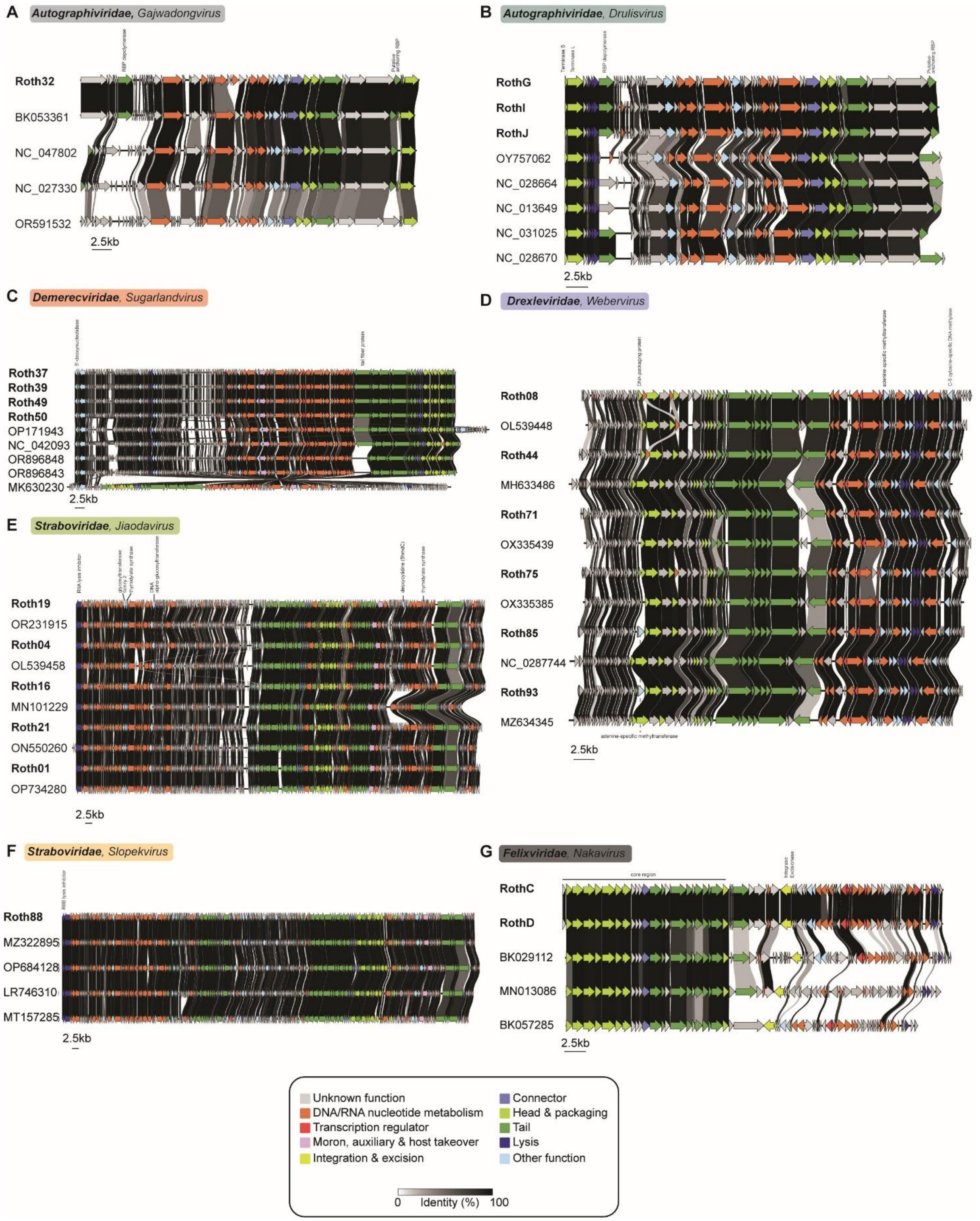


**Supplementary Figure S2.** Genome synteny plots of representative Roth phages with relatives. (**A-G**) Synteny plots were created by Clinker and genes were coloured by function. If necessary, reference genomes were re-zeroed to match the representative Roth phages genome organization for better comparison.

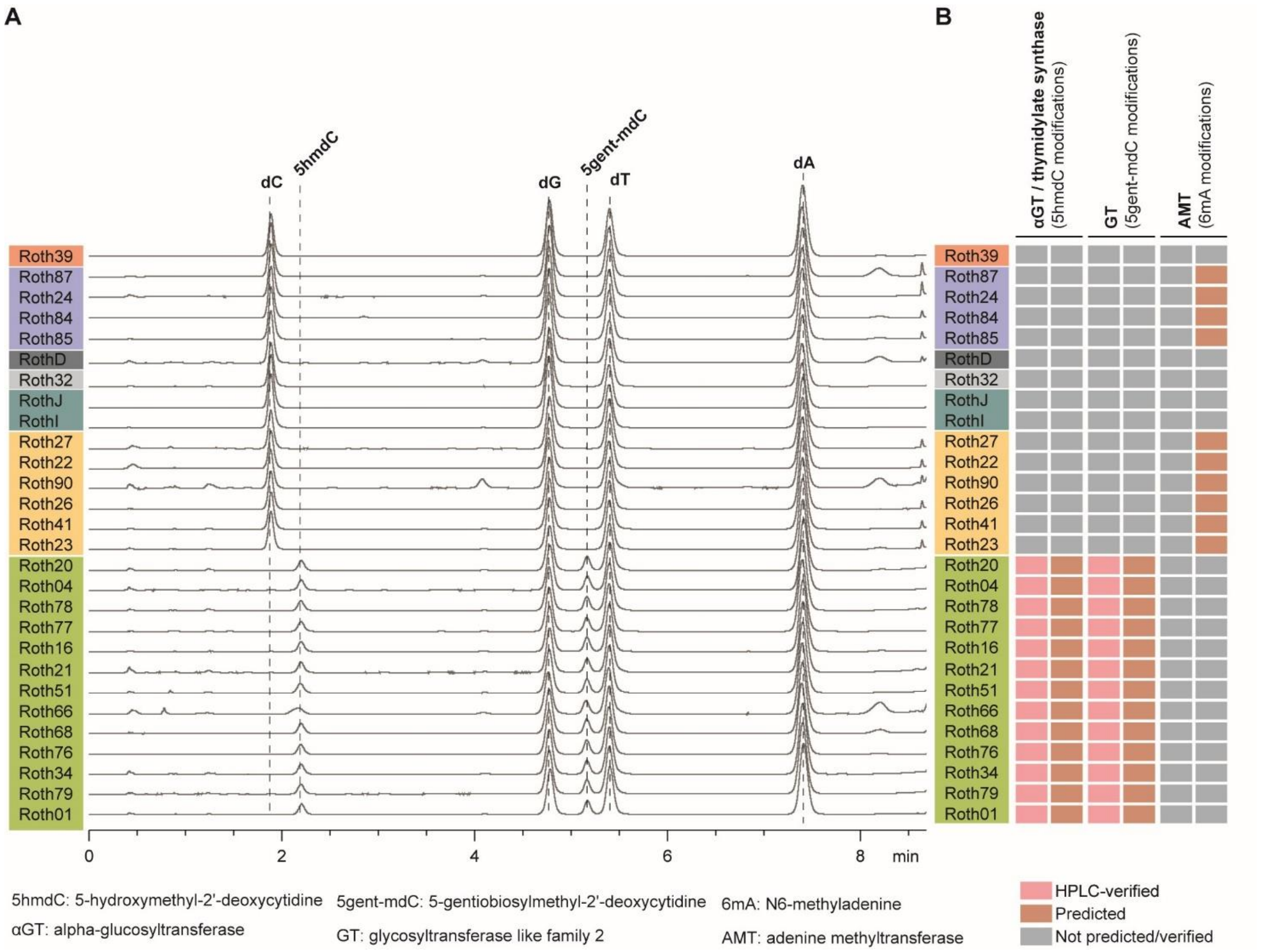


**Supplementary Figure S3.** DNA-modifying enzymes of Roth phages. (**A**) The DNA-encoded DNA modification enzymes were verified by HPLC for a random selection of phages covering all genera. (**B**) Summarised findings from HPLC in (A). Additional DNA-modifying enzymes were manually annotated and identified on their genomes.

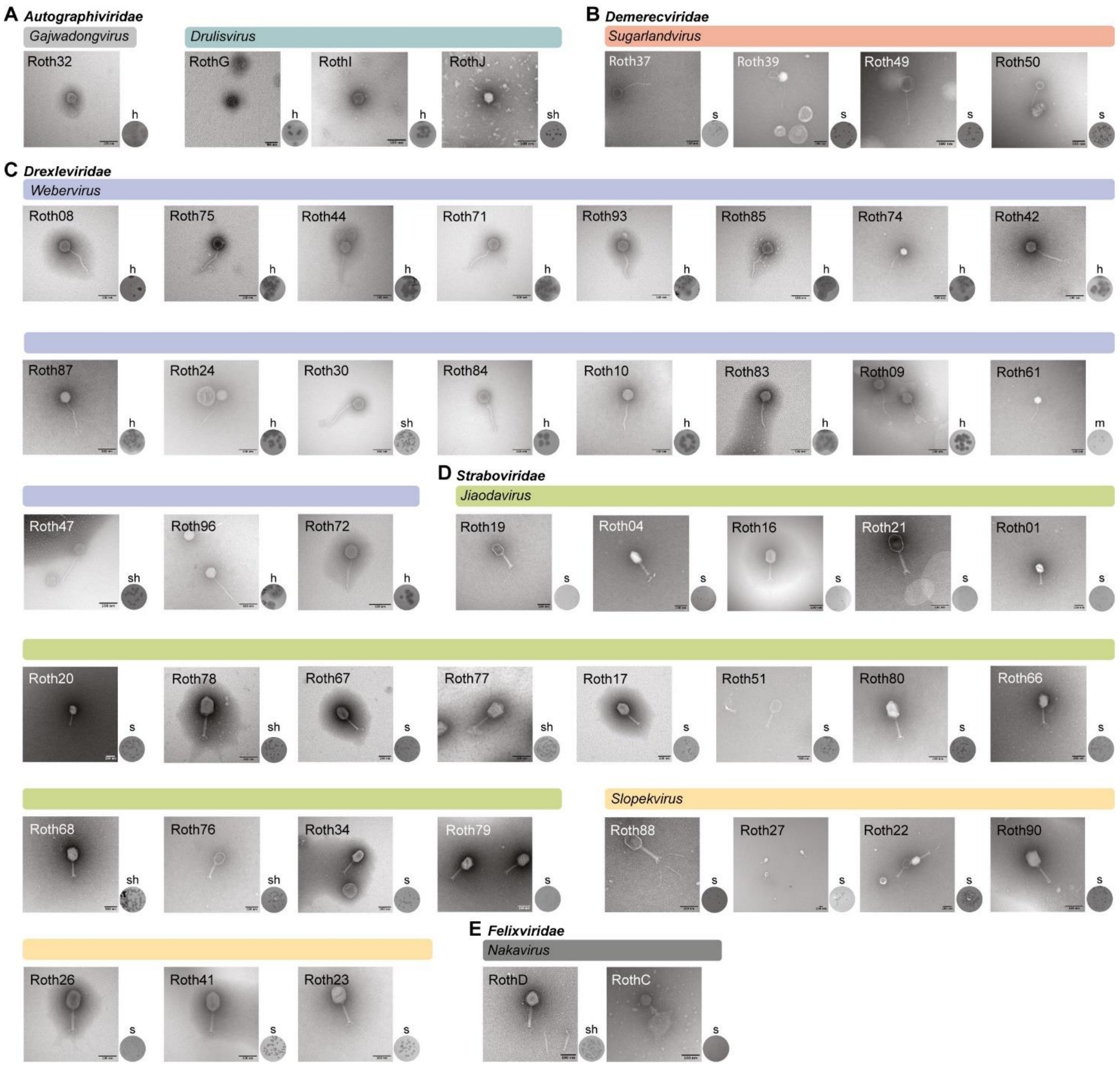


**Supplementary Figure S4.** Transmission electron microscope images of all KlebPhaCol Roth phages. (**A-E**) Bacteriophages were negatively stained with 5% Ammonium Molybdate (w/v) and 0.1-1% Trehalose, all bar scales are 100nm. Plaque morphology images are shown against the isolation hosts of each phage – except Roth49 and Roth50, which were plated against NCTC7427. Plaque morphology is summarised by a single letter above each image, s = small, m= medium, h = halo.

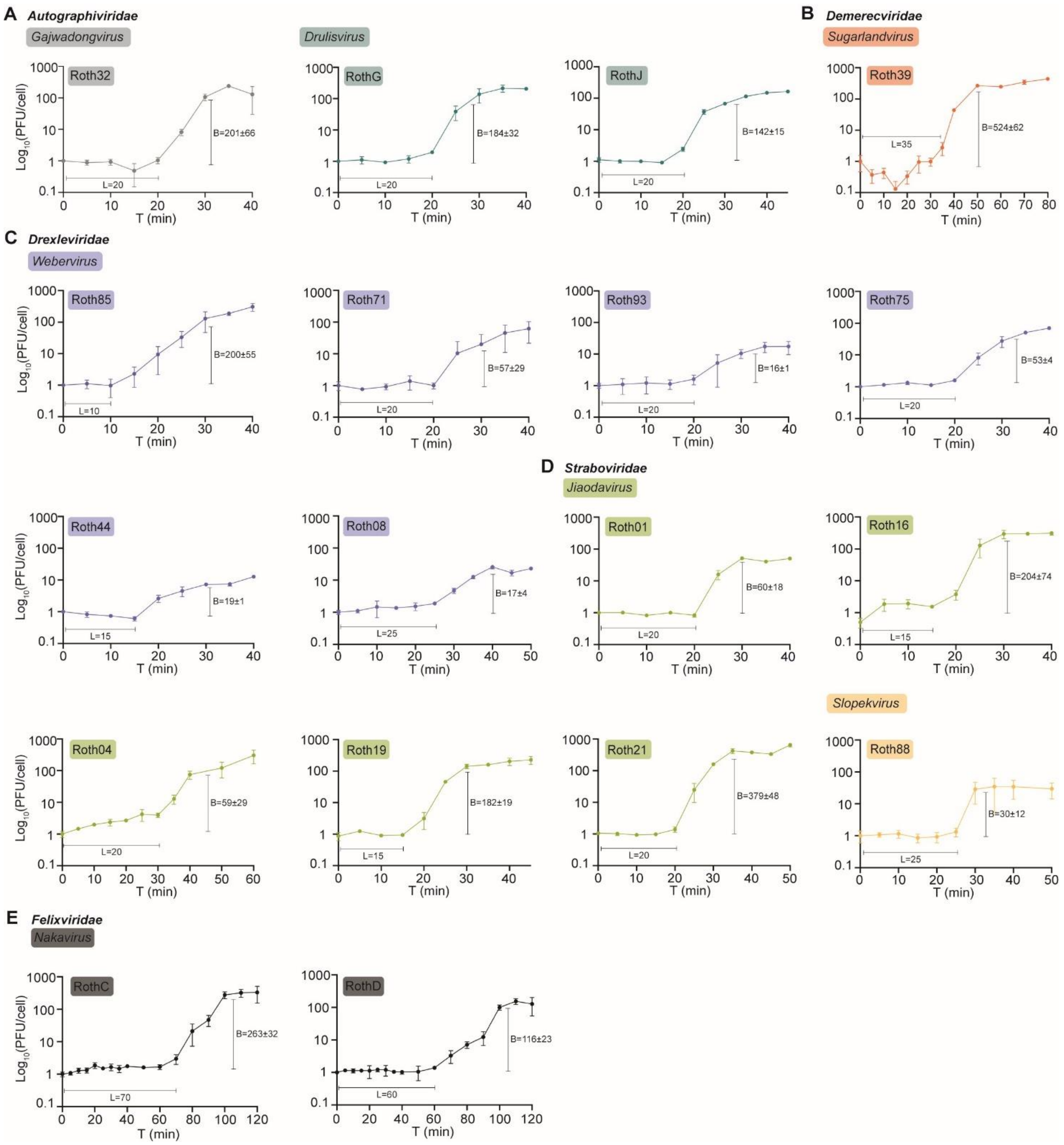


**Supplementary Figure S5.** One-step growth curves and burst size of all representative phages. (**A-E**) One-step growth curves were done in triplicate for each phage. Average ± standard deviation of burst sizes (B), and latent period (L) for each curve are also indicated.

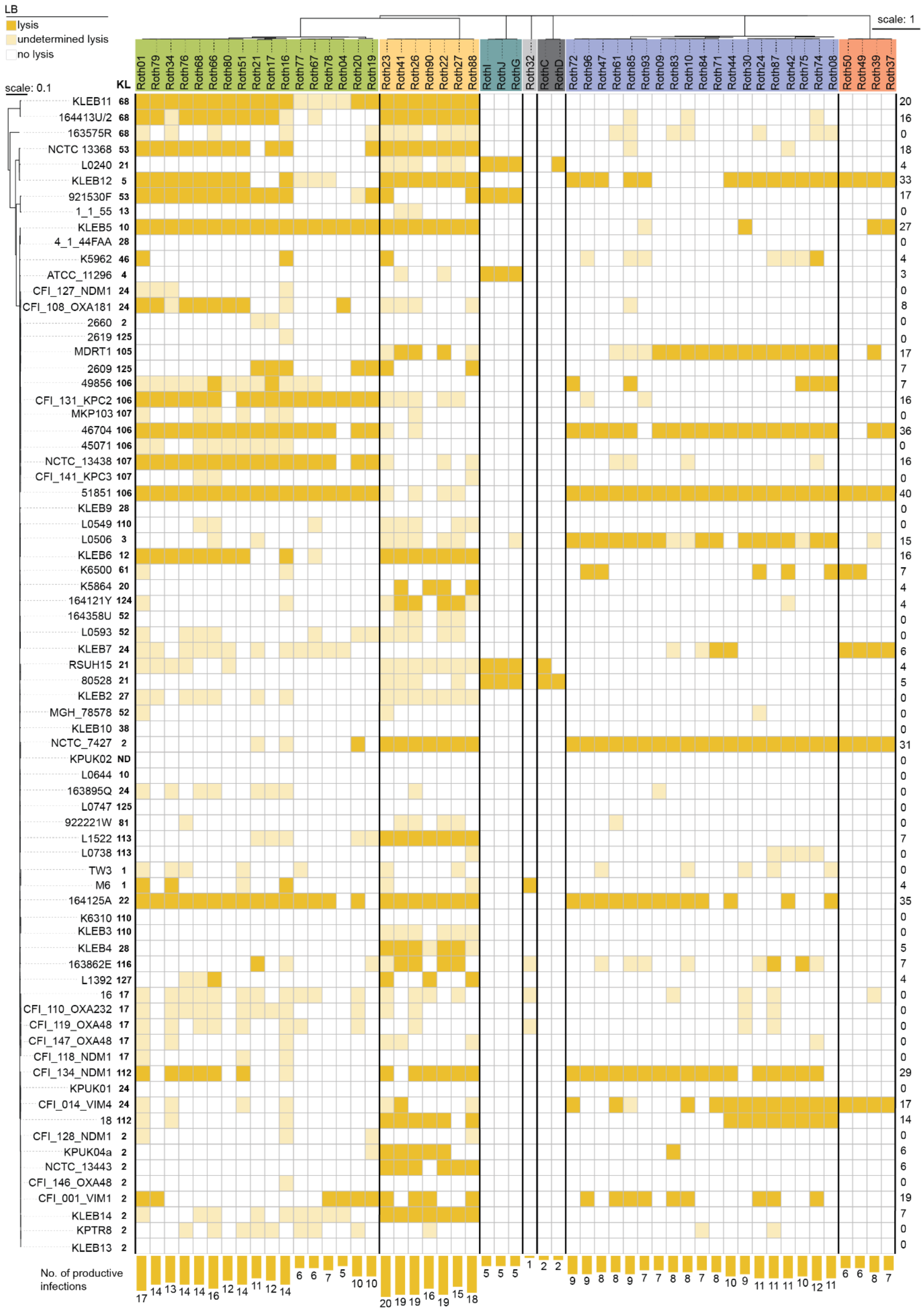

**Supplementary Figure S6.** Complete phage host range in LB medium. Heatmap is ordered by phage and strain phylogeny; phage phylogeny is divided by phage genus indicated by the colour blocks. Strain KL-type is displayed next to the strain name. Total counts of productive infections per strain are given on the right, and counts per phage are quantified and plotted as bars below the heatmap.

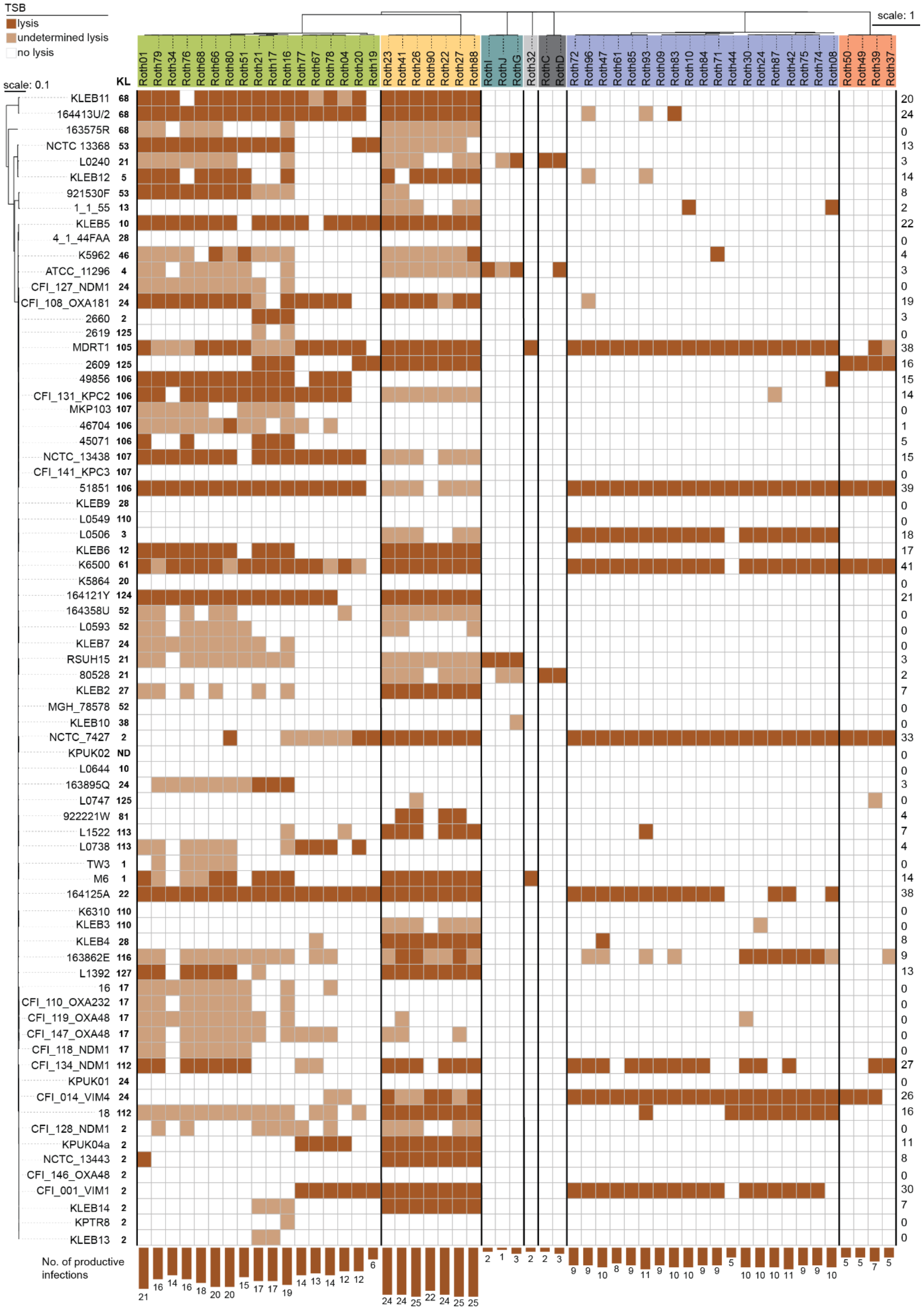

**Supplementary Figure S7.** Complete phage host range in TSB medium. Heatmap is ordered by phage and strain phylogeny; phage phylogeny is divided by phage genus indicated by the colour blocks. Strain KL-type is displayed next to the strain name. Total counts of productive infections per strain are given on the right, and counts per phage are quantified and plotted as bars below the heatmap.

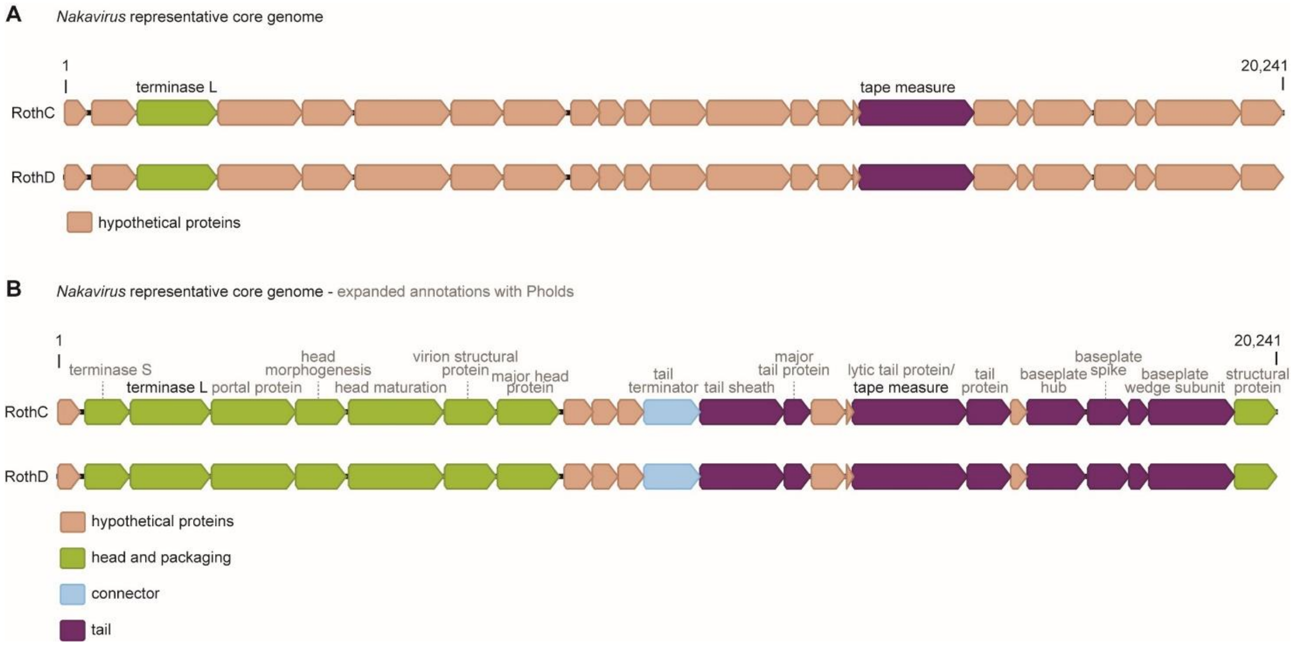


**Supplementary Figure S8.** Core genome annotations of RothC and RothD. (**A**) Represents annotations of the core region gp1 – gp23 (1-20,241bp) following a conservative annotation approach using sequence-based homology via MultiPhate annotation tool. (**B**) As (A) but following a structure-based homology annotation approach via Phold annotation tool, elucidating more functional genes.

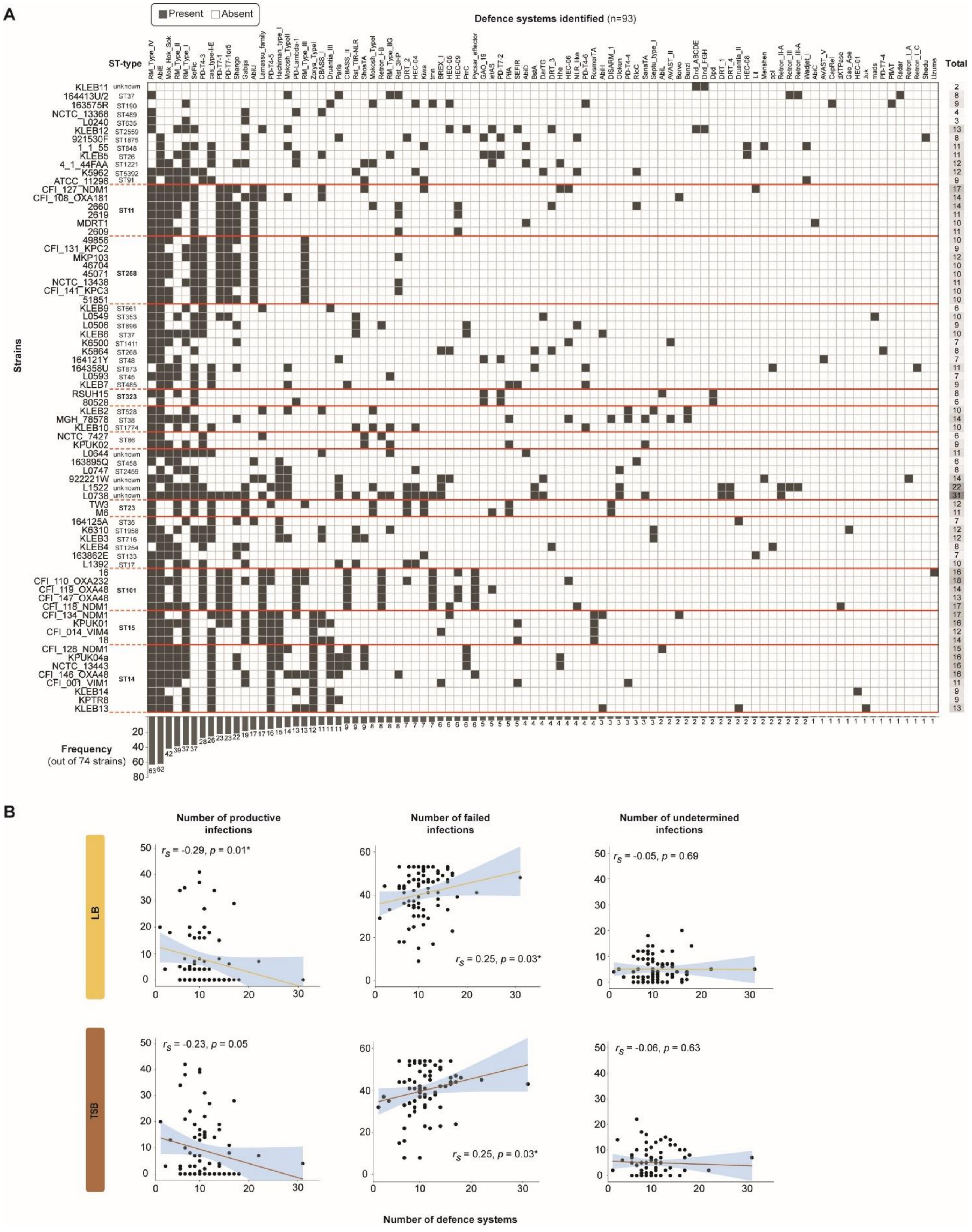


**Supplementary Figure S9.** Bacteria-encoded defence systems and correlation analyses with infectivity patterns. (**A**) Defence systems predicted by PADLOC and Defensefinder, showing only those experimentally verified as defence systems (heatmap is not showing PDC and VSPR systems, full table can be found in Supplementary Table S2)**.** Total count of defence systems encoded by each strain in shown on the far right. Frequency of the defence system in all 74 strains is indicated in the bottom of the heatmap. Strains are phylogenetically ordered, ST-type is indicated and grouped by the red lines. (**B**) Spearman rank correlation analysis was conducted and plotted using Rstudio ggplot2 to assess the relationship between the infectivity patterns of each strain and the number of encoded defence systems, considering only those systems that have been experimentally verified as showed in (A).

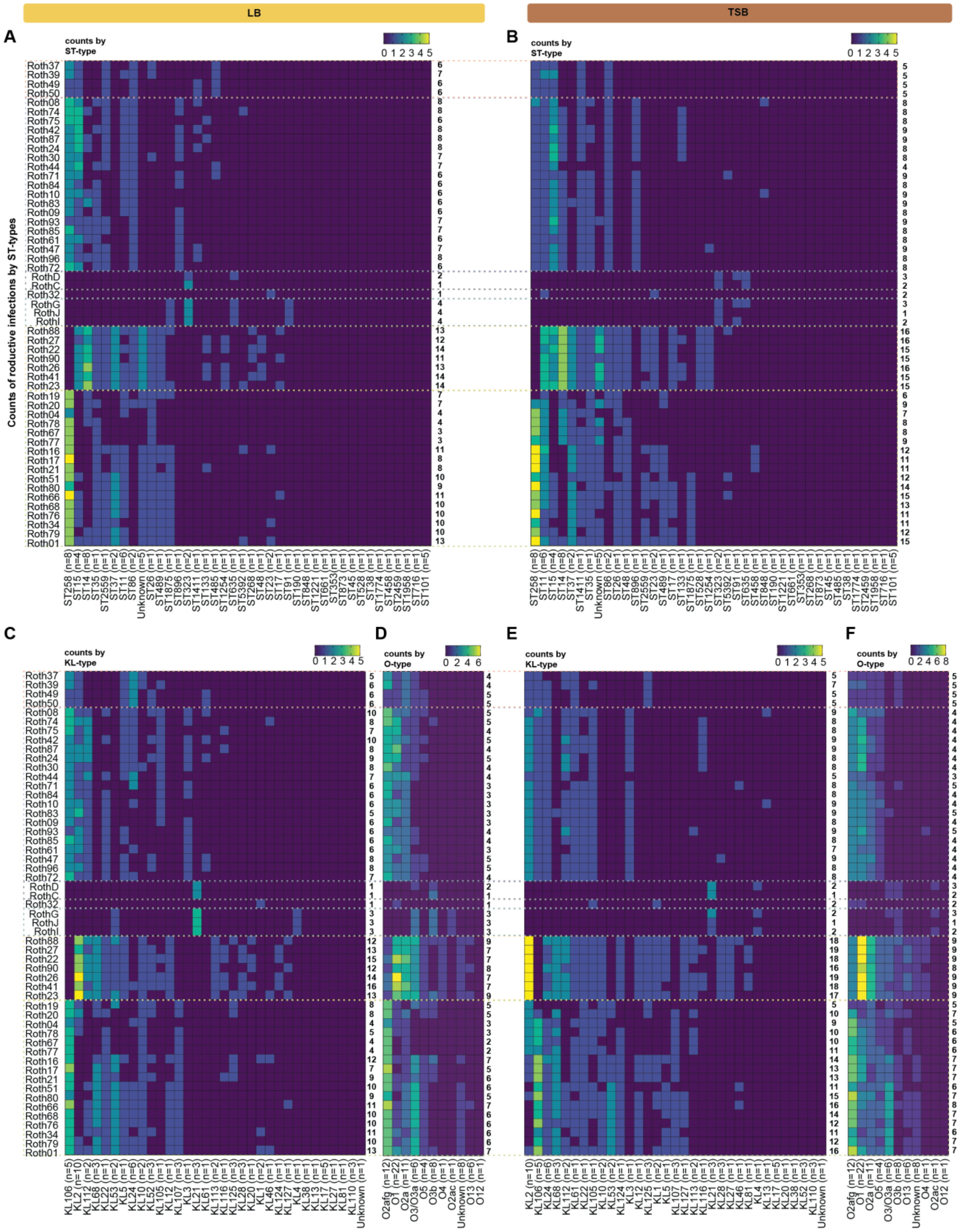


**Supplementary Figure S10.** Roth phage host range in terms of ST- KL- and O-antigen types. (**A**) Counts of ST-types infected per phage in LB medium. Total counts per phage are given at the right of the heatmap. Phages are ordered phylogenetically while ST-types are ordered in decreasing order of susceptibility. ST-types are followed by the total number of strains in the collection with that ST-type. (**B**) Same as a) but for TSB medium.

(**C**) Counts of KL-types infected per phage, and (**D**) O-antigen types infected per phage in LB. Phage and KL-type distribution follow the same logic as in (A). (**E,F**) Same as (C) and (D) but in TSB medium.

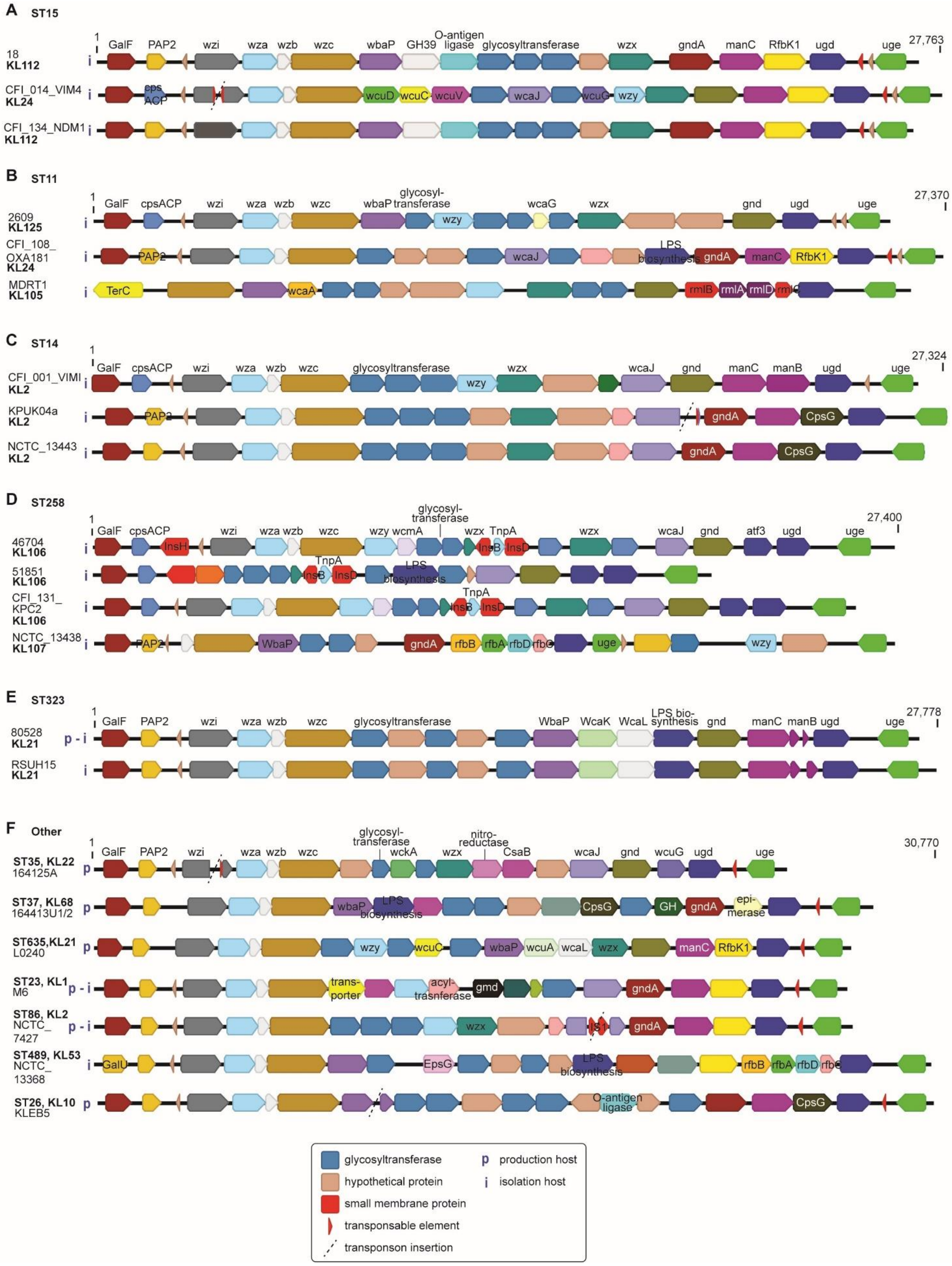


**Supplementary Figure S11.** Assembled capsule loci of isolation and production host strains. (**A-E**) Capsule loci are grouped by sequence-types (ST) with capsule locus (KL) types indicated below each strain name. (**F**)

Capsule loci grouped by host strains that do not share a ST-type with any other host. Isolation hosts are marked by an *i* while production hosts are marked by a *p*.

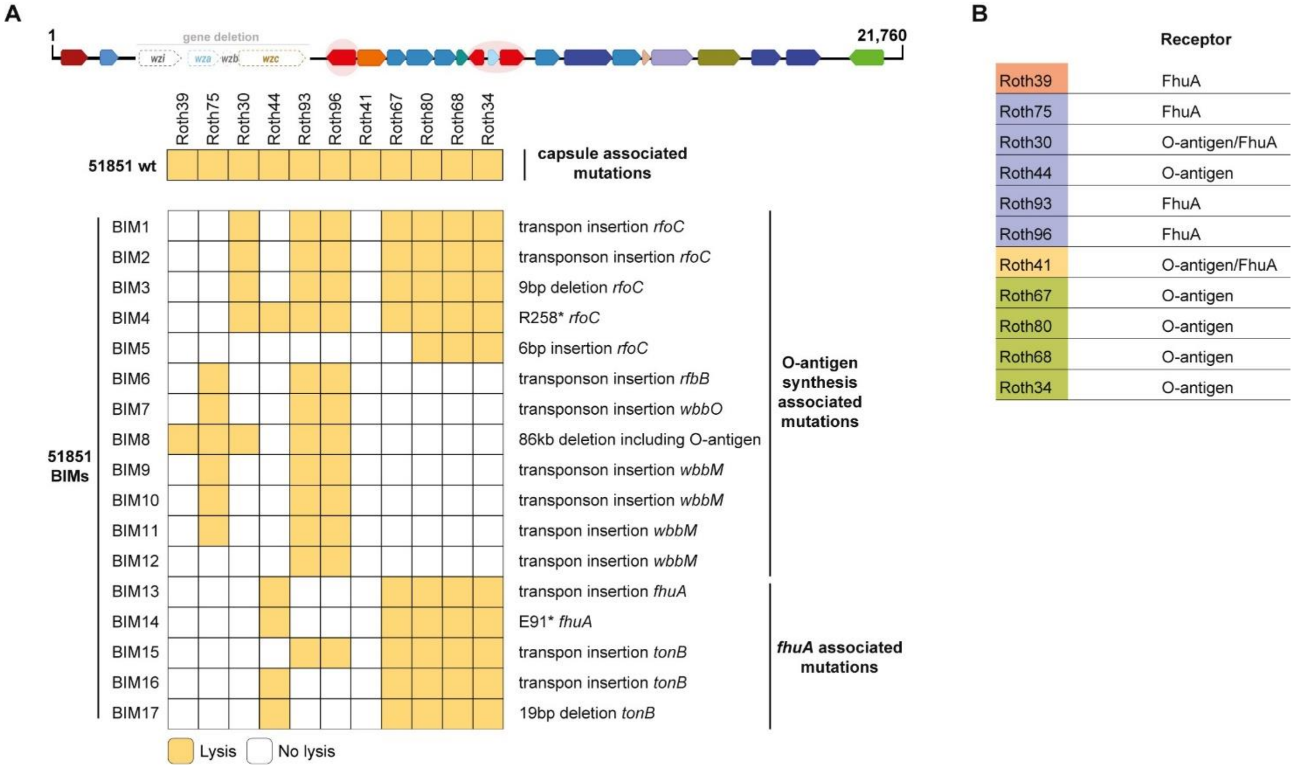


**Supplementary Figure S12.** Bacteriophage insensitive mutants (BIMs) reveal the receptors of selected phages. (**A**) Infectivity of the selected phages against the wildtype (wt) capsule-deficient strain 51851 and its BIM clones. Sequencing results revealed the BIM-encoded mutations to be in O-antigen synthesis associated genes or in the outer membrane receptor, *fhuA*. (**B**) Interpretation of results from (A) indicating likely receptors for the selected phages.

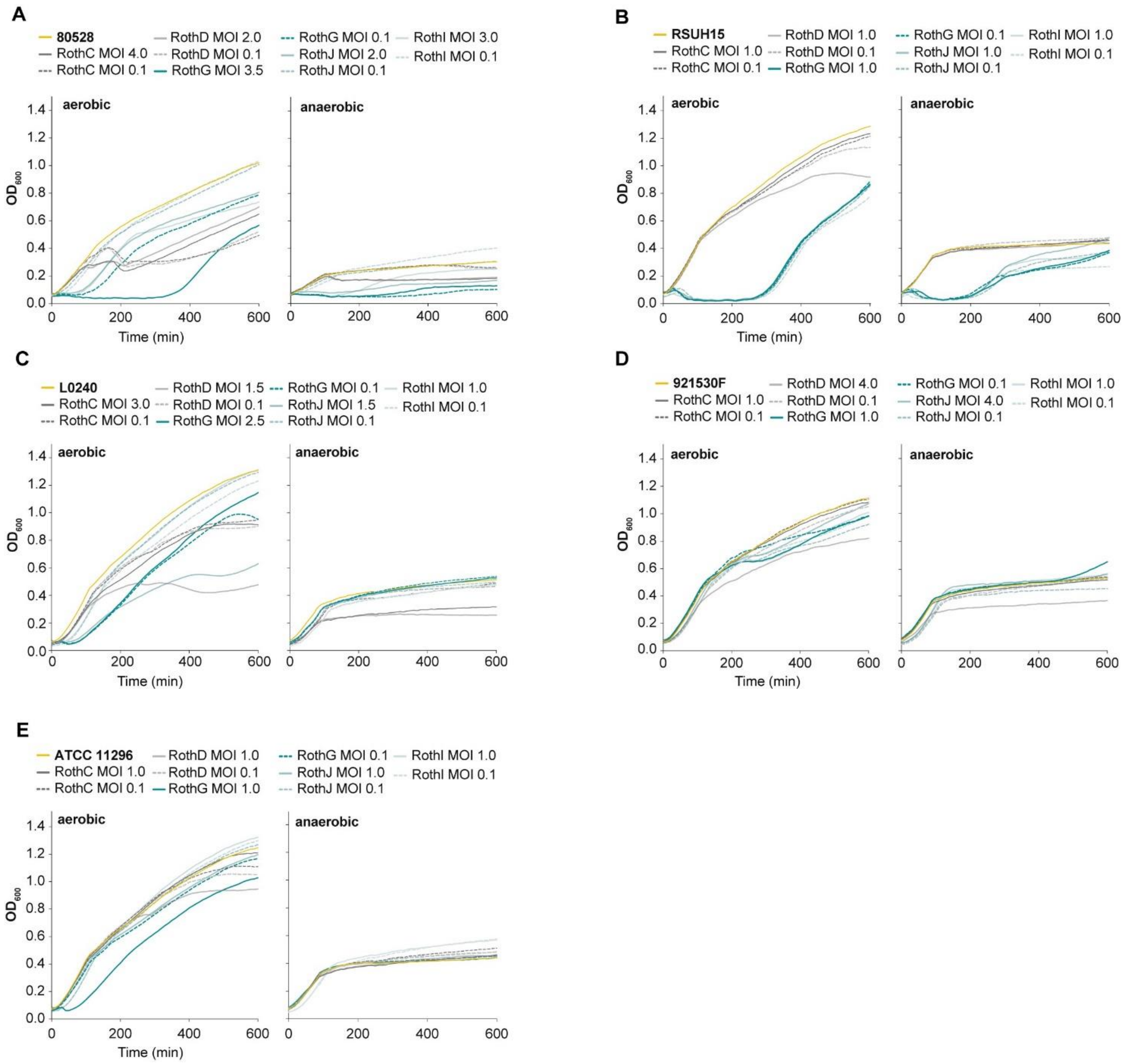


**Supplementary Figure S13.** Growth curves of ST323-targeting phages in susceptible strains. (**A-E**) Curves are the mean of three biological repeats. Bacterial control is shown in yellow, lower multiplicity of infection (MOI) curves are indicated by dashed lines.

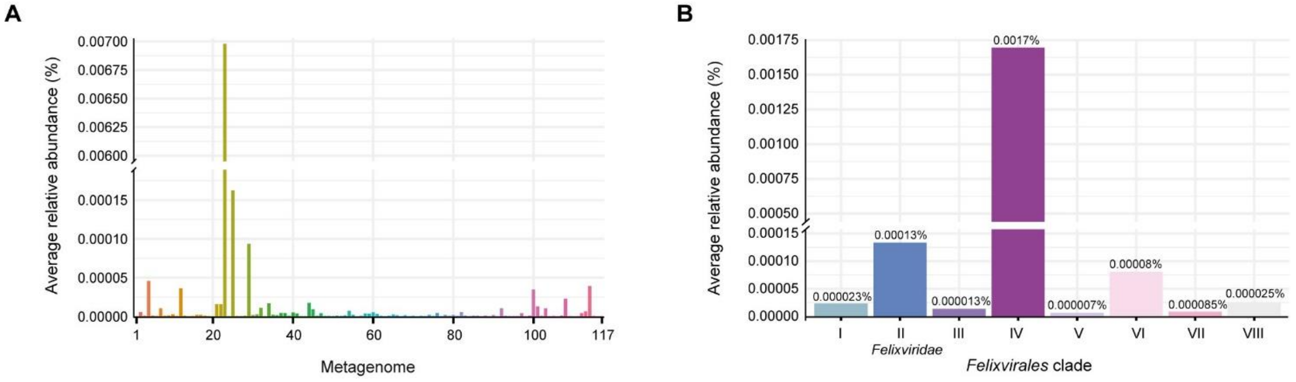


**Supplementary Figure S14.** Relative abundance of *Felixvirales.* (**A**) Relative abundance of reads per metagenome (117 total healthy adult metagenomes). (**B**) Relative abundance distributed by clade.
